# Cryo-EM Structure of Adeno-associated virus-4 at 2.2 Å resolution

**DOI:** 10.1101/2022.01.12.476110

**Authors:** Grant M. Zane, Mark A. Silveria, Nancy L. Meyer, Tommi A. White, Michael S. Chapman

## Abstract

Adeno-associated virus (AAV) is the vector of choice for several approved gene therapy treatments and is the basis for many ongoing clinical trials. Various strains of AAV exist (referred to as serotypes), each with their own transfection characteristics. Here, we present a high-resolution cryo-electron microscopy structure (2.2 Å) for AAV serotype 4 (AAV4). The receptor responsible for transduction of the AAV4 clade of AAV viruses (including AAV11, 12 and rh32.33) is unknown. Other AAVs interact with the same cell receptor, Adeno-associated virus receptor (AAVR), in one of two different ways. AAV5-like viruses interact exclusively with the <u>p</u>olycystic <u>k</u>idney <u>d</u>isease-like [PKD]-1 domain of AAVR while most other AAVs interact primarily with the PKD2 domain. A comparison of the present AAV4 structure with prior corresponding structures of AAV5, AAV2 and AAV1 in complex with AAVR, provides a foundation for understanding why the AAV4-like clade is unable to interact with either PKD1 or PKD2. The conformation of the AAV4 capsid in variable regions I, III, IV and V on the viral surface appears to be sufficiently different from AAV2 to ablate binding with PKD2. Differences between AAV4 and AAV5 in variable region VII appear sufficient to exclude binding with PKD1.

## 1. Introduction

Adeno-associated virus (AAV) is an icosahedral, non-enveloped, single-stranded DNA virus, which can infect humans. AAV was initially touted as a potential vector for gene therapy, because it is genetically amenable and the wild-type virus had not been associated with any diseases [1]. It is currently the gene-therapy delivery vector of choice in treatments for a number of monogenic genetic diseases such as Spinal Muscular Atrophy (SMA) [2,3]. When used as a vector to deliver a gene therapy, the genome of the virus is replaced with a gene of interest resulting in a recombinant AAV (rAAV). After some debate, consensus is that the risk of inducing hepatocellular carcinoma is minimal in those without liver disease [4,5] and no other disease has been connected with AAV infection, but a concern remains that particularly high vector loads of rAAV have been linked to fatal immunotoxicity in some test subjects [6,7]. Therefore, research to explore how various AAV strains (also referred to as serotypes) interact with cell receptors will hopefully add to our understanding of serotype-specific phenotypes and lay a foundation for improvement in the efficiency and specificity of vectors in different targeting modalities.

The AAV genome codes for three capsid proteins (VP1, VP2 and VP3). The DNA coding for the capsid proteins contains three different start codons such that all transcripts share the same C-terminal sequence of the VP3 protein (consisting of ∼540 amino acids) [8]. VP1 contains an ∼195 amino-acid extension beyond the N-terminus of VP3 while VP2 starts ∼60 amino acids upstream of the N-terminus of VP3. The ∼60 amino-acid sequence, found in both VP1 and VP2, is referred to as the VP1/2 common sequence. The estimated ratio of the three VP proteins in a mature capsid is 1:1:10. The portion of VP1 that is unique to VP1 (∼135 amino acids) is referred to as the VP1 unique region (VP1u). VP1u contains a phospholipase A_2_ (PLA_2_) domain [9-11], along with a putative nuclear-localization signal (NLS) [12,13]. In a mature capsid, VP1u and the VP1/2 common region are situated inside the capsid (with the genomic ssDNA) in locations that have not been apparent by X-ray crystallography and hinted in the cryo-electron microscopy of some AAVs, but not others [14-17]. It is not unexpected that the VP1u and VP1/2 regions have been refractory to high resolution structure, because they are present in only 10% and 20% (respectively) of the capsid subunits that must be 60-fold symmetry-averaged in most approaches to structure determination. Not only would the signal be much weaker than for fully symmetric parts, but they may be in a disordered state. During trafficking of the virus from the surface of a cell to the nucleus, VP1u and the VP1/2 common region become exposed and have been shown to be important for efficient infection [13,15,18-21].

Briefly, AAV infection occurs through the following steps. First, the virus binds to serotype-specific glycan attachment factors such as heparin sulfate or sialic acid on the surface of the cell [22], [23]. Next, the virus interacts with a protein surface receptor (AAVR for all serotypes except AAV4-clade members) to facilitate entry of the virus into an endosome [24]. In the endosome, the pH drops which results in a destabilization of the virus and VP1u is exuded [19]. Concomitant with the change in pH in the endosome, the calcium concentration also rises with the assistance of SPCA1, an ATP-powered calcium pump coded by the host’s ATP2C1 gene [21]. *In vitro*, VP1u can be exposed by heat treatment [18], but it is not yet clear what combination of lower pH, lower calcium concentration and other factors are the physiological trigger [19]. The presence of host protein GPR108 enhances transduction by AAV (except for AAV5), downstream of AAVR-binding and presumably during trafficking or uncoating [25]. Chimeric mutants show that GPR108-independence is conferred by the AAV5 VP1u, suggesting that, as VP1u is sequestered inside the capsid until late in endosomal trafficking, the GPR108 interaction may be important in endosomal escape. The virus is then trafficked to the nucleus with the assistance of the NLS regions located on the VP1/2 common region [13] where the viral genome (or, for rAAV, the transgene) is transported into the nucleus and expressed.

Differential patterns of molecular interactions between the various serotypes and AAVR are emerging. AAVR is a C-terminally membrane-anchored receptor with an ectodomain consisting of a signal peptide, a MANEC domain (motif at amino terminus with eight cysteines) and five non-identical polycystic-kidney disease domains (PKD) [26]. The tandem PKD domains are named PKD1 through PKD5. Initial studies of AAVR [27] showed that AAV2 was dependent primarily upon PKD2 for cell transduction, secondarily on PKD1 (in an accessory role), but not dependent on other PKD domains. AAV5 required PKD1 but not any other PKD domains, including PKD2. To follow up on these findings, cryo-EM images were taken of AAV2 in complex with either all five PKD domains (PKD15) [28] or with only the first two domains (PKD12) [29]. In both studies, AAV2 interacts tightly and specifically with the PKD2 domain with other domains of AAVR too flexibly oriented to be resolved. A similar pattern was seen with AAV1 when imaged in complex with PKD15 [30]. By contrast, cryo-EM images of AAV5 in complex with PKD15 [30] or PKD12 [31] show a complementary picture of a strongly bound PKD1, with PKD2 unresolved. Furthermore, AAV5 interacts specifically with the PKD1 domain at a viral site that is distinct from that by which AAV2 binds to PKD2. Overlay of the AAV2 and AAV5 complexes show that it would not be possible to connect the observed PKD1 and PKD2 domains into a single polypeptide in a hypothetical unseen AAV2 complex with the PKD1 domain oriented as in the AAV5 complex [31].

In the current work, we focus on AAV4 since it is a serotype that is not dependent on AAVR for successful cellular transduction. Cryo-EM is used to extend the resolution of the AAV4 structure beyond that of the prior crystallographic structure at 3.2 Å (PDBid: 2g8g) [32]. The crystal structure had been a seminal advance in 2006, because, with just 59% sequence identity to the then only high resolution structure, the type-species AAV2 [33], it represented the diversity among human AAVs. It provided an opportunity to define surface segments of the primary sequence as variable regions in structure (VR-I through VR-IX), that would, in time, become associated with distinctive functional phenotypes of AAV serotypes. The AAV4 crystal structure had been preceded by a cryo-EM structure [34] at 13 Å resolution, impressive in 2005, but then sufficient only to dock in an AAV2-based *pseudo*-atomic model. A “cryo-EM resolution revolution”, based on improved microscope stability, fast and sensitive detection devices and improved computational data processing [35-37], has made possible the determination of AAV structure at higher resolution than by crystallography [38,39]. Fourier image theory tells us that the information content of a 3D structure at 2.2 Å resolution is 3-fold higher than at 3.2 Å. Thus, with the goal of understanding how inter-serotype structural differences might underpin an ability to use AAVR for cellular entry, the need was met for an AAV4 structure of precision comparable to those of AAVR-binding serotypes [28,31]. With structures available for several AAV4 homologs (AAVrh32.33, 11 and 12 [40,41]) we can explore the features, conserved among AAVR-independent serotypes, that are significantly different compared to AAV-binding serotypes.

With the assumption that the interactions between AAV and AAVR are conserved within closely related serotypes, we sought to determine which features of AAV are important for binding to AAVR. For the PKD2 interface, structures of receptor complexes were available for AAV2 (and AAV1) [28-30], and we could also overlay structures of uncomplexed AAV capsids, known or presumed to interact analogously [27,42]. For the PKD1 interface, receptor complexes were available for AAV5 [30,31] and homologous serotype structures could similarly be overlaid. With a high resolution AAV4 structure in-hand, and related structures overlaid, we then consider how differences between the AAV4 clade and corresponding parts of other clades might impact AAVR interactions. There are differences between AAV4 and AAV5 that plausibly explain why AAV4 does not interact with AAVR through PKD1, and differences between AAV4 and AAV2 that rationalize the lack of interactions mediated through PKD2.

## 2. Materials and Methods

### 2.1 Expression and purification of AAV4

Virus-like particles (VLPs) for AAV4 were expressed in *Spodoptera frugiperda* Sf9 cells from a polyhedron H promoter (P_*polH*_) on a bacmid. The bacmid was constructed in *E. coli* DH10Bac cells following transformation with the pFastBac LIC cloning vector (4A) (Addgene #30111) containing a cloned copy of the AAV4 VP1 gene. The VP1 gene was cloned into the destination plasmid using ligation-independent cloning (see Table S1) [43]. The start codon for VP1 was mutated to ACG in order to down-regulate expression of VP1 so as to accommodate a desired ratio of the three capsid proteins (1:1:10), as was done elsewhere [31,44]. Sequencing of the cloned gene was performed at the University of Missouri DNA core facility and compared with the published sequence of AAV4 (accession number NC_001829). Following expression in Sf9, the VLPs were isolated and purified as described previously [45] except that purification by heparin-affinity, expected to work only for serotypes with heparan sulfate attachment factors, was replaced by additional iterations of cesium chloride gradient ultra-centrifugation [31]. AAV4 VLP was then dialyzed into HN buffer (25 mM HEPES, 150 mM sodium chloride, pH 7.4) and used in additional experiments.

### 2.2 Single Particle Cryo-EM sample preparation and imaging

Samples of dialyzed AAV4 VLP (2 µL at 0.33 mg/ml) were placed upon glow-discharged EM grids (copper grids with lacey carbon, Ted Pella cat. no. 01824) and allowed to adhere for 2 min. Using an FEI Vitrobot Mark IV, the sample was blotted (blot force: 4; time: 2 s; temperature: 25 °C; and humidity: 100%) and plunged into liquid ethane. Cryo-EM images were recorded at the Pacific Northwest Cryo-EM Center with a Gatan K3 detector and Gatan BioQuantum energy filter on a Titan Krios 3 electron microscope at 300 keV. A data set was collected at high-magnification (4809 images collected at 165,000x) as 80-frame movies at 0.256 Å/pixel at a total dose of 48 electrons/Å^2^/movie. Defocus varied between -0.6 and -1.8 µm.

### 2.3 Image processing

The Cryo-EM images were processed using Relion 3.1.1 [46]. The movie files were loaded and motion corrected using the built-in (RELION’s own) implementation. CTF estimation was performed in RELION on the motion corrected images with CTFFIND-4.1 [47]. Particles were initially picked in RELION with LoG-picker and sorted by 2D classification to establish a template which was used by the auto-picker to generate a larger set of particles. This expanded set of particles was then sorted into 20 2D-classes. The particles from the 18 best 2D classes were extracted, refined and used to do a 3D reconstruction where I1 icosahedral symmetry operators were applied. Particles then underwent per particle CTF estimation and per particle motion correction and refined a second time. After a final masking step a coulombic potential map of 2.2 Å was obtained.

### 2.4 Atomic Modeling and Refinement

RSRef [48] was used to calibrate the magnification of the AAV4 Coulombic potential map against the 3.2 Å structure of AAV4 obtained from X-ray crystallography (PDB: 2g8g, [32]). The magnification was adjusted by 0.41%. The published AAV4 structure was overlaid on the adjusted AAV4 electron density map; backbone and rotamer adjustments were performed manually within Coot 0.8.9.2 [49]. This was followed by using RSRef-embedded CNS [48,50] to perform stereochemically-restrained, all-atom real-space refinement (parameterized in Cartesian space). The final correlation of the model to the experimental 3D Coulombic potential map was 0.86 for all map grid points within 2.4 Å of any non-hydrogen atom.

### 2.4 Thermal shift assay

Thermal shift assays were performed with VLPs of serotypes AAV2, 4 and 5, and these were compared with one another. Each 20 µL sample contained 0.125 mg VLP/ml (HEPES-buffered), 1x sample dye and 1x supplied buffer (Protein Thermal Shift Dye Kit, Thermo Fisher 446-1146). Each sample was run in triplicate. The samples were prepared on ice and allowed to incubate in the dark for 20 minutes before being subjected to thermal denaturation on a QS3 thermocycler (Applied Biosystems). The samples were incubated at the designated temperature (20°C to 99.9°C) for 1 min, read for fluorescence, heated by 0.3°C and repeated. The data were processed using a local interface to a custom Thermal Shift Assay script (https://beamerlab.shinyapps.io/software/ “TSA (1-X, many-Y)”) based upon the calculations described elsewhere [51].

### 2.5 ELISA binding assays

ELISA experiments were performed as previously described [31]. Two ELISA experiments were conducted: one to test for binding of PKD1-2 to AAV4 and another to show that the AAV4-specific antibody (ADK4), but not the AAV5-specific antibody (ADK5b), could bind to AAV4 VLP. To check for binding of PKD12 to AAV4, various AAV serotypes (AAV2, 4 and 5) were adhered to a 96-well ELISA plate (Costar, 9018), bovine serum albumin (BSA; ThermoFisher, BP9703-100) was used to block the plate, a sample of 6xHis-tagged PKD12 was allowed to bind to the virus and an HRP-conjugated antibody (ProteinTech, 66005) allowed to bind to the His-tagged ligand.

AAV4 (and AAV5) VLP preparations were validated using the serotype-specific monoclonal antibodies ADK4 (Progen, 610147) and ADK5b (Origene, AM09121PU-N). VLPs were adhered to the ELISA plate, the wells were blocked with BSA, the primary mouse antibody was allowed to bind to AAV, then secondary antibody added (HRP-conjugated, goat derived; ThermoFisher, 62-6520). Following binding of the HRP-conjugated antibody, the HRP substrate (Abcam, 171523) was added, then hydrochloric acid (ThermoFisher, A144C-212) to stop the reaction. The signal was read at 450 nm (BioTek, Synergy H1). All samples were run in triplicate.

### 2.6 Structures analyzed

The structures for the homologous AAV structures were obtained from the PDB. For AAV1 bound to PKD15, the structure PDBid 6jcq [30] was used. For AAV2 bound to PKD15, the structure 6nz0 [28] was used. For AAV5 bound to PKD12, the structure 7kpn [31] was used. For AAV4, the structure 7thr, determined in this study, was used. For AAV7, the structure 7l5q [40] was used. For AAV8, the structure 2qa0 [52] was used. For AAV9, the structure 3ux1 [53] was used. For AAV11, the structure 7l6f [40] was used. For AAV12, the structure 7l6b [40] was used. For AAVrh32.33, the structure 4iov [41] was used. The phylogenetic tree was constructed with ClustalX [54].

## 3. Results and Discussion

### 3.1 Biophysical and biochemical confirmation of AAV4

AAV4 VLPs were validated by comparison to the expected characteristics: thermal denaturation (thermal shift assay, Table S 2) and recognition by specific antibodies (ELISA). As expected, the AAV4 VLP reacted with monoclonal ADK4 and not to the AAV5-specific antibody, ADK5b (data not shown).

### 3.2 AAV4 is not bound by AAVR

Transduction data for AAV4 show no dependence upon AAVR [42], which suggests that AAV4 might not bind to AAVR. This could also merely reflect the presence of a dominant alternative means of entry. Virus overlay assays had previously shown no evidence of binding between AAV4 and fragments of AAVR (PKD1, PKD2, PKD3 or PKD15) that included all plausible binding domains [42]. However, these negative results were treated with an abundance of caution, because viral overlay assay assumes that interactions remain possible in the environment of a denaturing gel, and this is not always true. Thus, AAVR-AAV4 binding was additionally measured by ELISA using the PKD12 construct (containing the first two PKD domains), for which the positive results with AAV2 and AAV5 provide a robust control (Figure S 1). Again, AAV4 does not appear to interact with AAVR.

### 3.3 High-resolution structure of native AAV4

The prepared AAV4 VLP was imaged by electron cryo-microscopy (cryoEM) and the reconstruction was refined to 2.21 Å, according to the Gold standard Fourier shell correlation curve (Figure S 2). The quality of the 3D reconstruction is shown in Figure 1.

**Figure 1:**
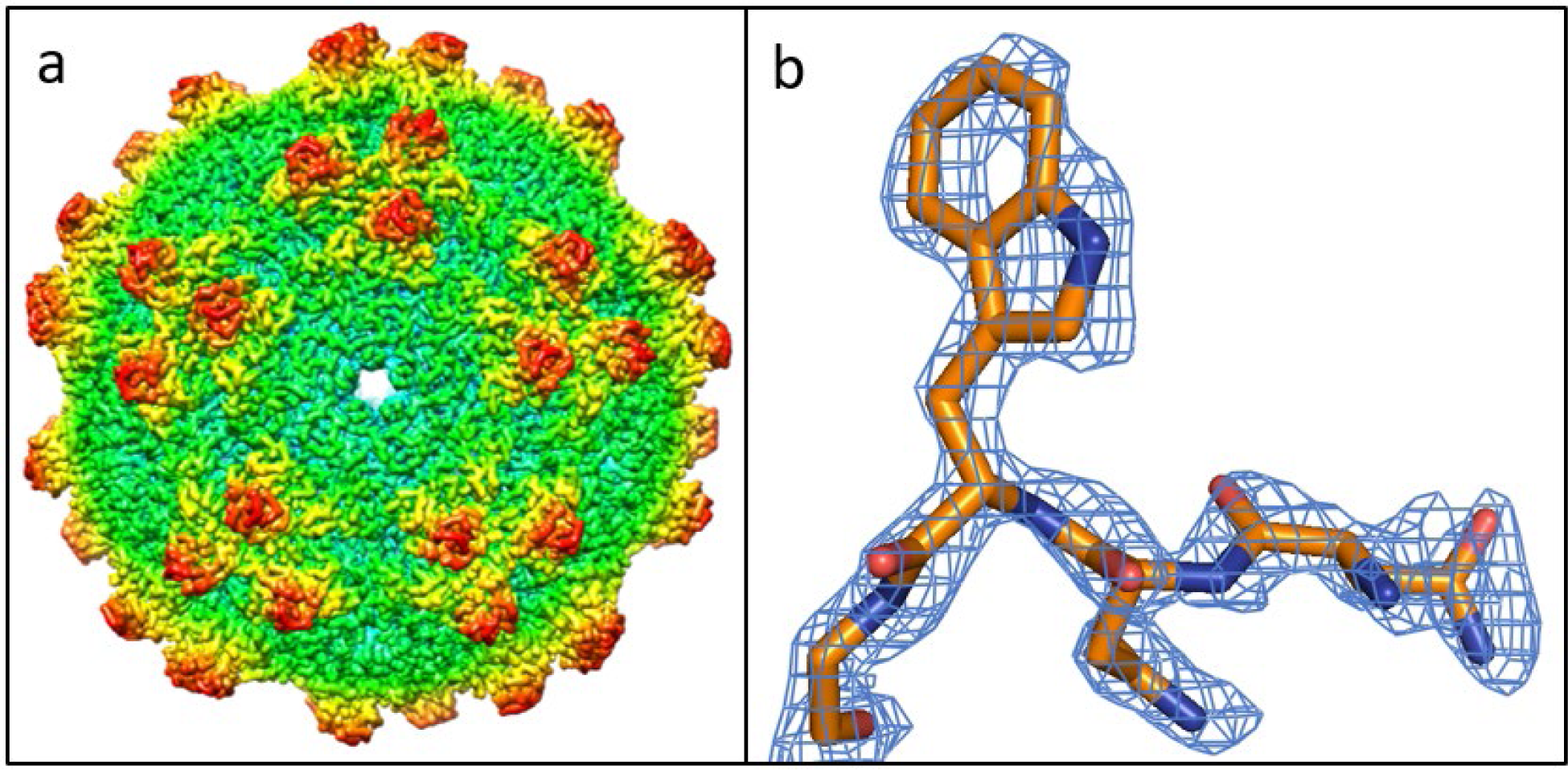
(a) Cryo-EM reconstruction of AAV4 at 2.21 Å resolution, color-coded by distance from the center. The view is down one of 12 pores running along 5-fold axes of symmetry. Groups of 3 spikes, related by 3-fold axes of symmetry encircle the 5-fold. (b) Coulombic potential for residues AAV4-T240 through -L243.

### 3.4 Overlay of AAV4 upon AAVR-bound AAV

We compared the current AAV4 structure with published AAVR-AAV structures to identify components of the AAV4 structure, at amino acid level, that render AAV4 incompatible with the binding of AAVR. The AAV4 structure was aligned to several AAVR-bound complexes using the align function within PyMol [55,56].

#### 3.4.1. Comparison of AAV4 at the PKD2-binding site as seen in AAV2 and AAV1

For AAV2, residues from VR-I, -III, -V, -VI, -VIII and -IX (plus a few others) are in contact with PKD2 [28,29] (Table 1). AAV4 and other serotypes were aligned to the structure of the AAV2-PKD12 complex. Each of the contact residues were examined for changes in steric hindrance electrostatic charge, or more subtle differences could impact the ability of AAV4 to bind to AAVR.

**Table 1:**
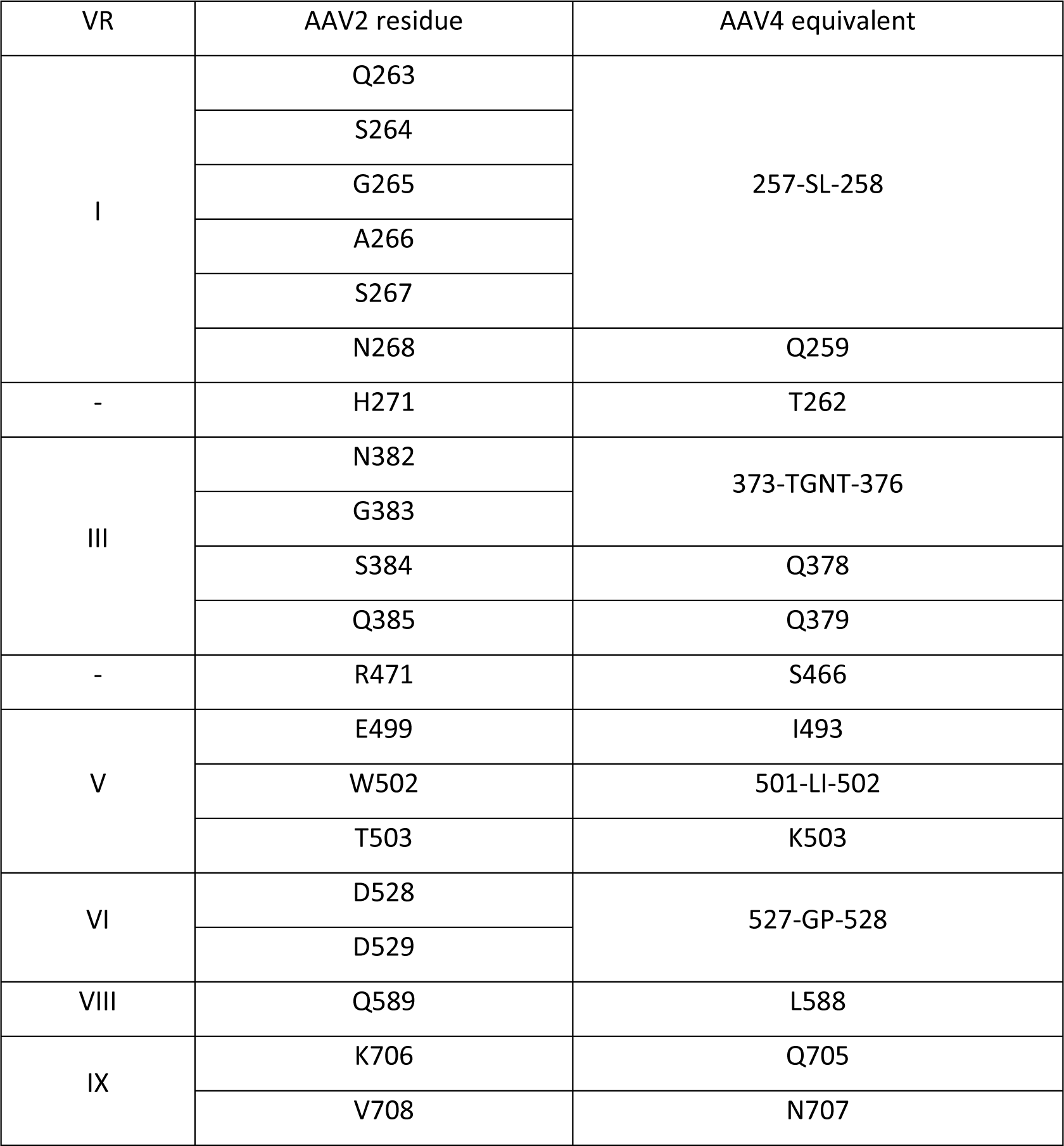
Residues on the surface of AAV4 that align with the residues of AAV2 that contact AAVR PKD2 [28].

VR-III contains four residues that are important for the binding of AAV2 to PKD2. There is considerable variation between AAV2 and some of the other serotypes (Figure 4). VR-III of AAV4-clade members has a structure that is distinct from that shared by the majority of AAVs that, like AAV2, are PKD2-binders (Figure 4). Within the AAV4-clade, AAV4 is the outlier in sequence and structure, but AAV4-clade members are much more similar to each other than to other AAVs. Steric conflict would be substantial between AAVR-K438 and AAV4-T376. Conflict at the backbone of AAVR-D437 is somewhat less, but significant, and there is also conflict predicted with a conserved asparagine in AAV11, 12 and rh32.33. AAV5, like AAV4 does not bind AAVR PKD2. Although AAV5 N375 and T376 are translated ∼2Å relative to AAV4, this same region of VR-III would clash with PKD2 near K438 in both AAV5 and the AAV4 clade (Figure 4).

**Figure 2:**
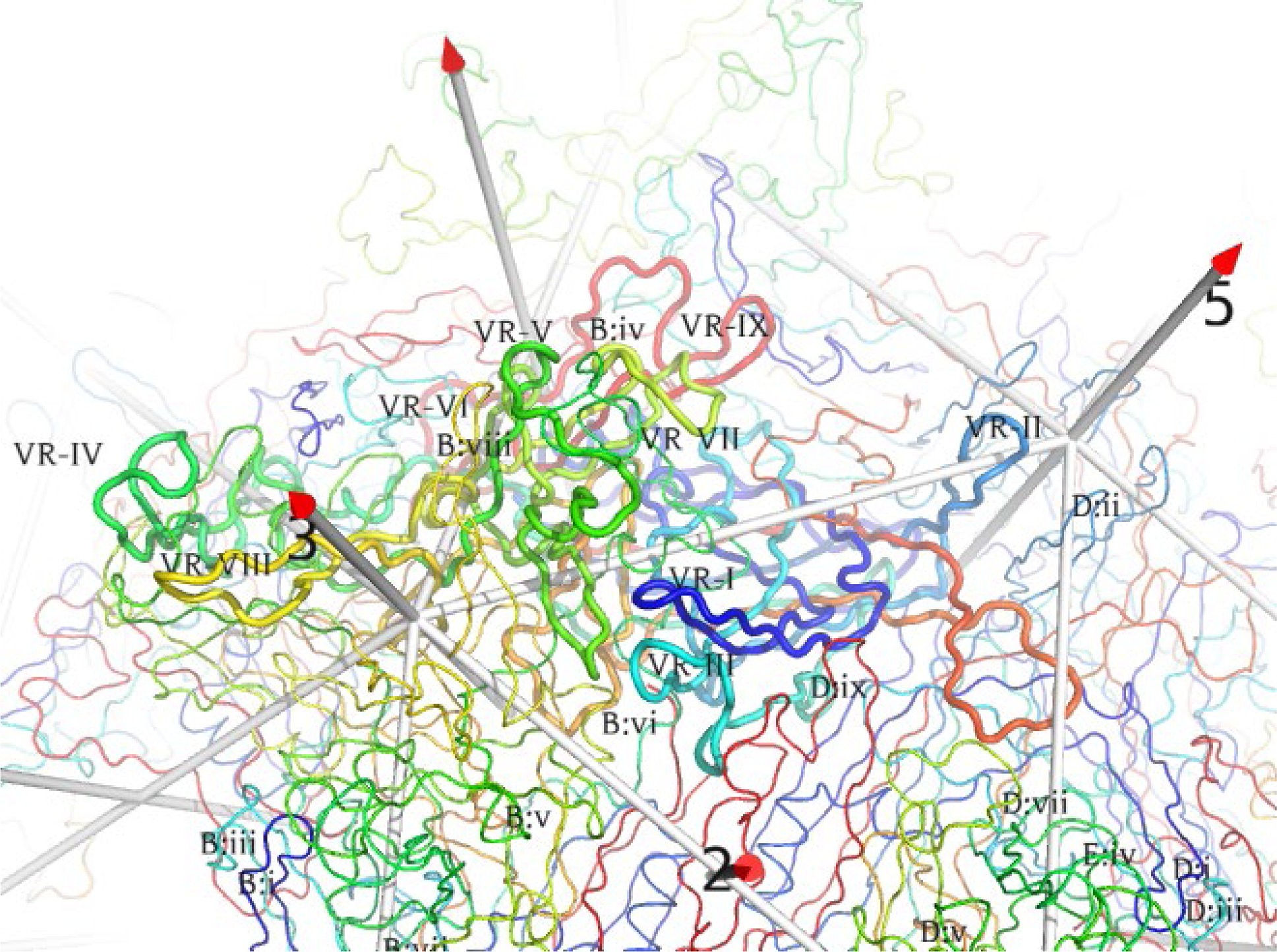
Juxtaposition of variable regions (VR) as the major capsid protein (VP3) is surrounded by symmetry neighbors (thinner trace). The view is from outside the capsid, looking down an icosahedral 2-fold, with a 3-fold left, and a 5-fold right. The conserved β-barrel is behind the outer surface loops that dominate the foreground. Each subunit is rainbow-colored from N-terminus (blue) to C (red). The AAV4 cryo-EM structure is annotated by sequence-variable region (VR) [32]. VR-IV through VR-VIII are all contributed by a long loop between β-strands G & H. VRs of those neighboring subunits most closely intertwined are annotated in abbreviated form (chain-id:loop#).

**Figure 3:**
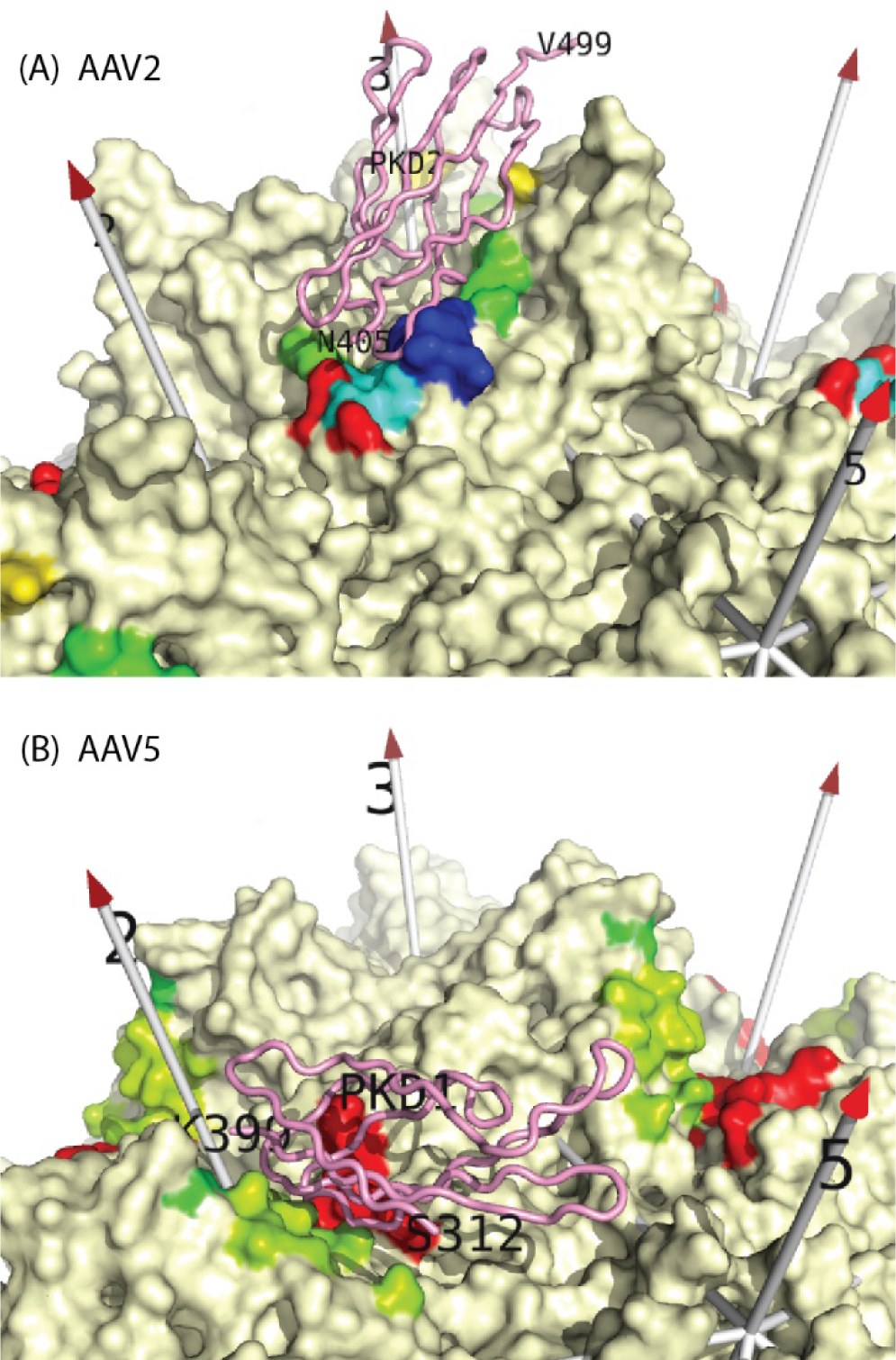
Binding sites for AAVR on AAV2 and AAV5. Single copies of the PKD2 and PKD1 domains of AAVR (pink backbone) are shown as they are bound in structures of AAV2 (top) and AAV5 (bottom) complexes respectively [28,31]. The viral surfaces are viewed with a similar perspective as Figure 2, using the same rainbow coloring, but only for amino acids that come within a 4.5 Å distance of receptor atoms (or their symmetry-equivalents). Comparison with Figure 2 highlights the parts of AAV4 that correspond to the receptor interfaces of AAV2 and AAV5.

**Figure 4.**
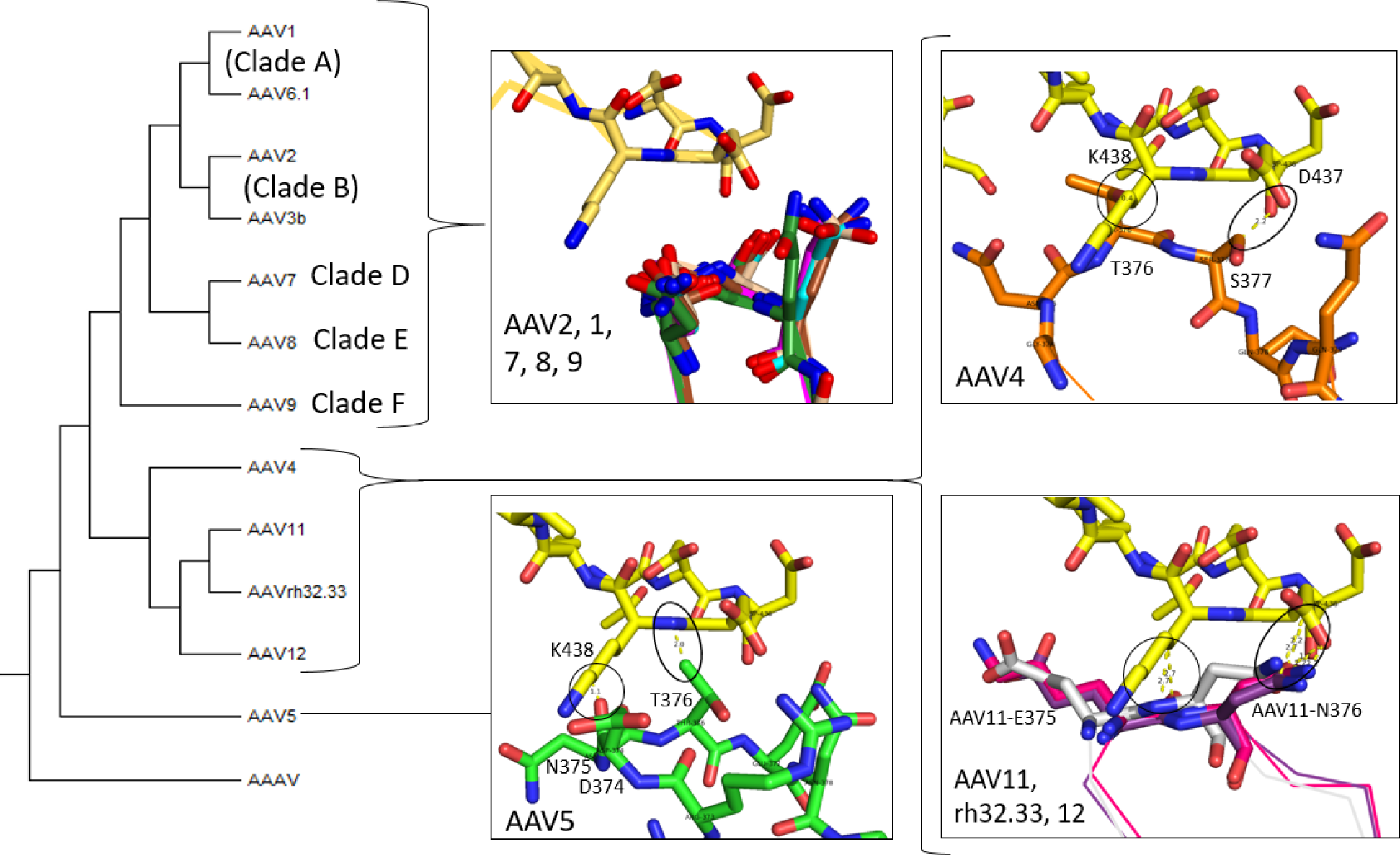
Potential interactions of variable region 3 (VR-III) for AAV structures after alignment to the AAV2-PKD15 receptor complex (PDBid: 6nz0). Carbons are colored yellow for AAVR and by serotype for AAV structures. The following structures were used: AAV1 (dark green; PDBid 6jcq), AAV2 (magenta; PDBid 6nz0), AAV4 (orange; PDBid 7thr), AAV5 (green; PDBid 7kpn), AAV7 (brown; PDBid 7l5q), AAV8 (wheat; PDBid 2qa0), AAV9 (cyan; PDBid 3ux1), AAV11 (purple; PDBid 7l6f), AAV12 (red; PDBid 7l6b), and AAVrh32.33 (grey; PDBid 4iov). There is steric hindrance between AAVR-K438 and AAV4-T376 (or AAV5 D374–N375 or AAV11, 12, rh32.33) (circles) and a second clash between the side-chains of AAVR-D437 and AAV4-S377 (asparagine in AAV11, 12 or rh32.33) (oval). The phylogenetic tree is based on VP1 sequences.

VR-I is immediately adjacent to VR-III in the 3D structure. In AAV2, six of the residues interacting with PKD2 come from VR-I with one additional residue (H271) immediate downstream. As shown in Figure 5, this region in AAV4 (and most other serotypes) shows important differences from that seen in AAV2. Surprisingly, steric clashes are seen in AAV7, 8, 9, where AAVR appears to clash with AAV7 and AAV9 at a single residue (S268 in both strains) and with multiple residues in AAV8. AAV5 clashes with PKD2 at two AAV5 residues: S254 and G257. AAV4 and AAV12 each contain two residues (AAV4 S257 & Q259 and AAV12 T266 & N268) that would clash with PKD2. AAV11 and AAVrh32.33 have a single residue (T257) that would conflicts with PKD2 (D436 & D437).

**Figure 5.**
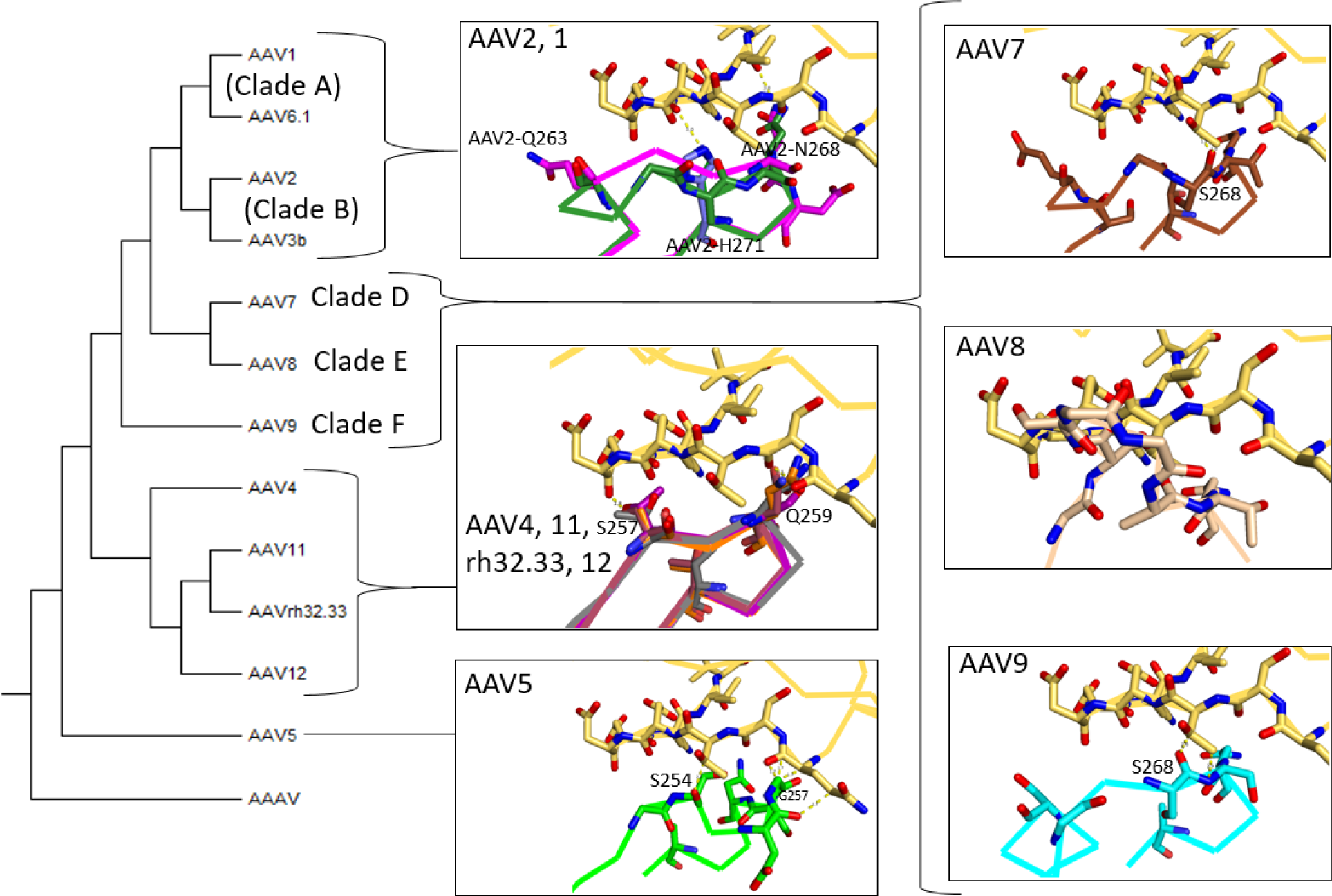
Potential interactions of variable region 1 (VR-I) for AAV structures after alignment to the AAV2-PKD15 receptor complex (PDBid: 6nz0). Structures and serotype color coding are as described in Figure 4, and AAVR is shown with yellow carbons. There are multiple clashes between PKD2 and most of the AAV serotypes, including the AAV4-clade and AAV5. Surprisingly, there appear to be clashes in many of the AAVR-dependent strains (AAV7, 8, 9), which are thought to interact with PKD2, suggesting that conformational adaptation may occur with some serotypes. The phylogenetic tree is based on VP1 sequences.

The third region of AAV2 that interacts with PKD2 is VR-V, which contains three residues of interest. Presented is a comparison of VR-V between various serotypes (Figure 6). All AAV4-clade members show a substantial clash between VR-V and PKD2 that is unique to the clade. It is noteworthy that although AAV5, like AAV4, does not bind PKD2, the structure of AAV5 VR-V is very similar to those of the AAV2-like serotypes that bind PKD2, so VR-V is not the cause of the AAVR binding differences between AAV5 and AAV2.

**Figure 6.**
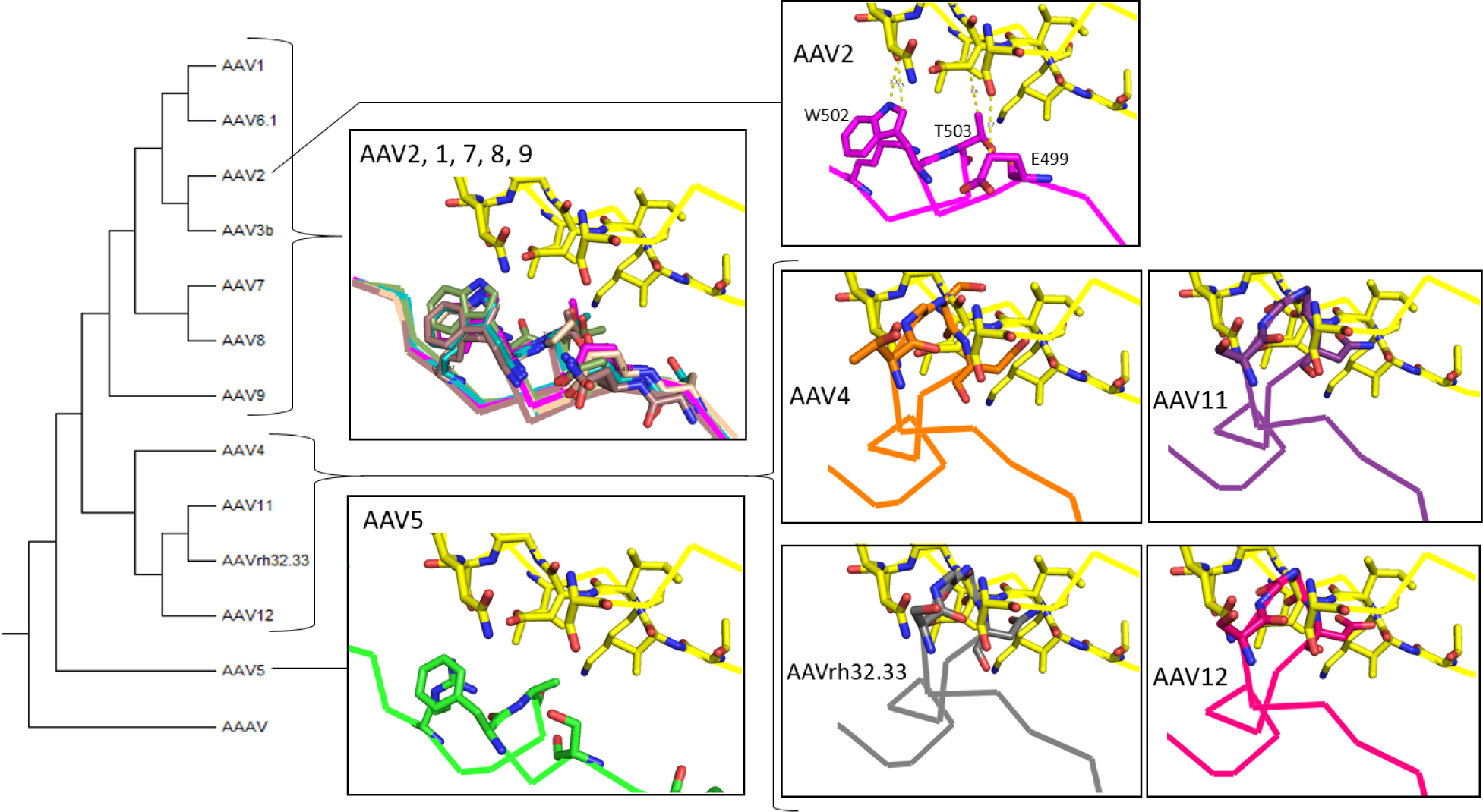
Potential interactions of variable region 5 (VR-V) for AAV structures after alignment to the AAV2-PKD15 receptor complex (PDBid: 6nz0). Structures and serotype color coding are as described in Figure 4, and AAVR is shown with yellow carbons. There would be considerable steric clashes between PKD2 and all AAV4-clade members for residues P494-Y504 in AAV4 and corresponding residues in other clade members. There are no clashes for known PKD2-binders like AAV2, but also no clashes for non-binding AAV5. The phylogenetic tree is based on VP1 sequences.

The next region of AAV2 that interacts with PKD2 is VR-VI, which contains two AAV2 residues in close contact. VR-VI from various serotypes is compared in Figure S 3. The loop in AAV4 clade members is further removed from where PKD2 would be and would likely not be part of any interaction.

PKD2 interacts with AAV2 VR-VIII from two symmetry-equivalent subunits. In the first, AAV2-Q589 contacts AAVR-S425 (Figure S 4, located at the beginning of PKD2-βB), an interaction that is conserved for AAV1 and AAV9, but not others. The second subunit makes contact through the positively charged AAV2 R588 and R585 of the heparin-binding domain (HBD) which interact with negatively charged AAVR E458 & D459, in the PKD2-βD-βE loop at distances of 2.7 Å and 3.2 Å, respectively (Figure 7). Interestingly, neither residue was identified by either of the AAV2-complex papers even though the binding residues in AAVR were identified [28]. All serotypes have polar amino acids in this location, but it is only clade B that has the arginines of the HBD capable of a salt bridge electrostatic attraction. (Each AAV2 capsid has 60 symmetry-equivalent RXXR, but at each one, the glycan attachment and PKD2-binding interactions would be mutually exclusive [28].) The length of the arginine side chains in clade B allows a favorable PKD2 electrostatic interaction without distortion of either the AAV or AAVR structure. Indeed, we do not see systematic differences in the docking of PKD2 between clade B (AAV2) and clade A (AAV1) [28,30]. We divided the VR-VIII interactions according to the two AAV subunits involved (Figure 7, Figure S 4), but neither shows any potential for steric hindrance from any of the superimposed AAV structures. The conclusion is that the favorable influence of the HBD in clade B is modest, and that there is no other reason to consider VR-VIII to be a major determinant in the mode of AAVR-binding.

**Figure 7.**
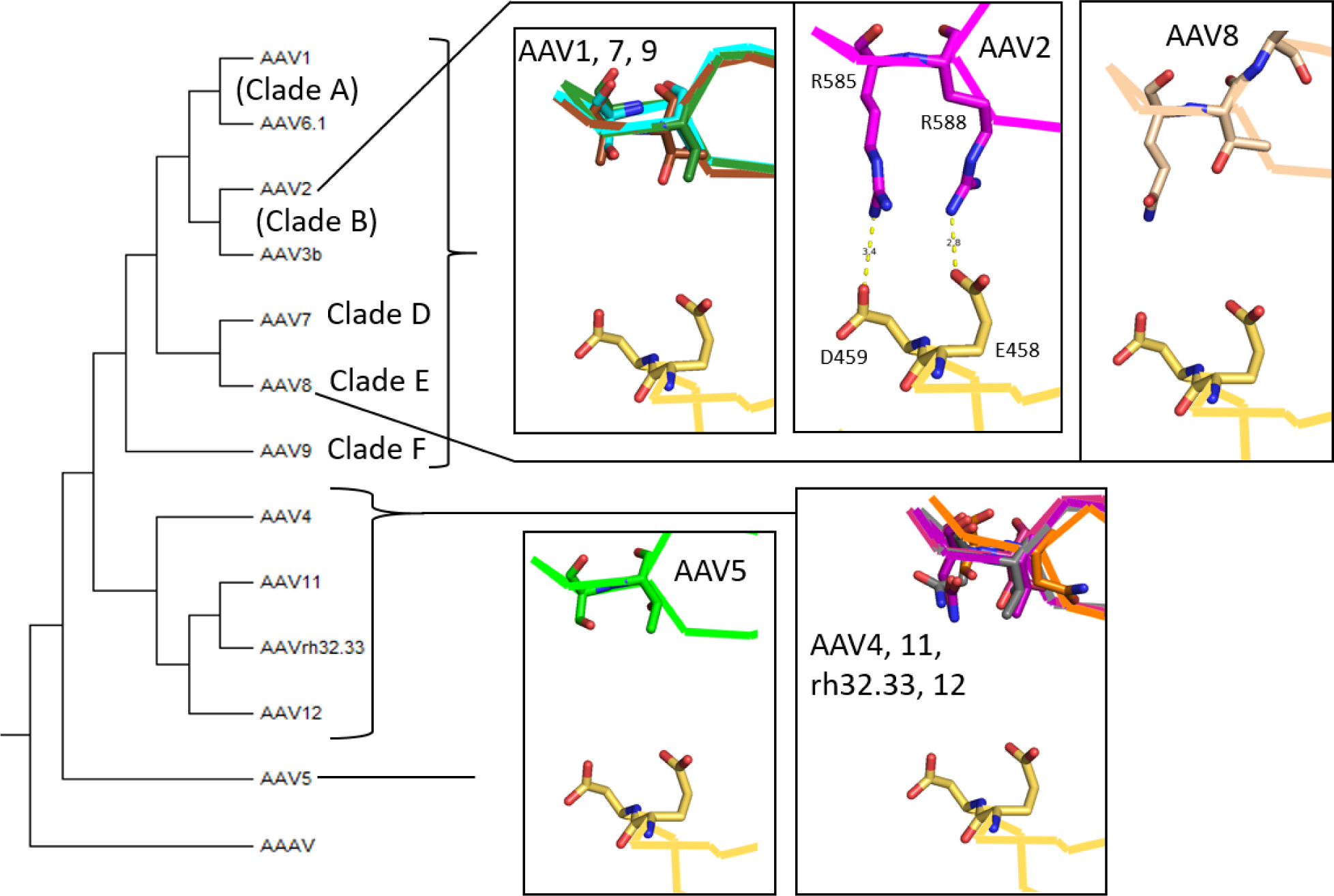
Potential interactions of variable region 8 (VR-VIII) for AAV structures after alignment to the AAV2-PKD15 receptor complex (PDBid: 6nz0). Structures and serotype color coding are as described in Figure 4, and AAVR is shown with yellow carbons. Interactions of a symmetry-equivalent VR-VIII with a different region of PKD2 are shown in Figure S 4. Neither of the AAV2 contact residues, R585 nor R588 (interacting with AAVR D459 and E458 respectively), are conserved outside clade B. None of the serotypes analyzed would conflict with PKD2 as bound in AAV2. The phylogenetic tree is based on VP1 sequences.

The final region to be considered with respect to PKD2 is VR-IX (Figure 8). Relevant to the observed 3.2 Å salt bridge between AAV2-K706 and AAVR-D437 [28], the lysine is conserved in all clades except AAV5 and AAV4-like. Replacing the lysine is an aspartic acid in AAV5 and one of several polar residues (asparagine, glutamine or threonine) in the AAV4-clade. In summary, from VR-IX we have a potentially favorable electrostatic contribution from all clades known to bind PKD2. This is neutralized in the AAV4 clade with a switch to a polar residue. In AAV5 (which binds PKD1, but not PKD2), the switch to a like (repulsive) charge is in the context of a shorter side chain increasing the distance and mitigating the incompatibility.

**Figure 8.**
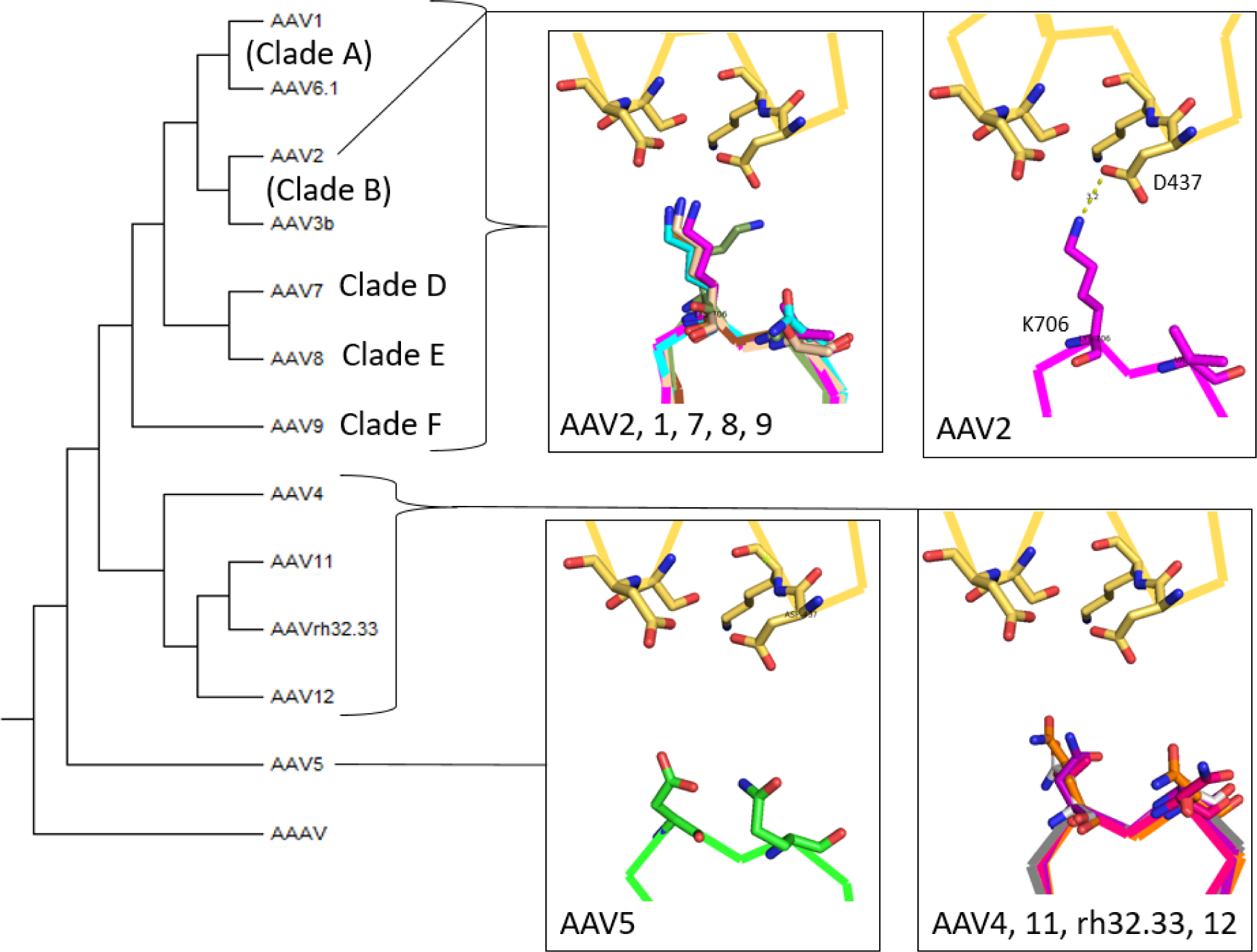
Potential interactions of variable region 9 (VR-IX) for AAV structures after alignment to the AAV2-PKD15 receptor complex (PDBid: 6nz0). Structures and serotype color coding are as described in Figure 4, and AAVR is shown with yellow carbons. There would be no steric hindrance for this region between PKD2 and these serotypes. However, the positive charge located at AAV2-K706 (and which is conserved in serotypes AAV1, 7, 8, 9) is not conserved within the AAV5 and AAV4-clade members. The phylogenetic tree is based on VP1 sequences.

Even though VR-II, -IV and -VII of AAV2 were not seen to interact with PKD2, these regions were analyzed in case any were sufficiently different in AAV4 to account for lack of binding between AAV4 and PKD2. VR-IV is located upstream of R471. A comparison between serotypes for this region is presented in Figure 9. The VR-IV loop in the AAV4 clade has a conformation that is different from other clades and incompatible with the binding of PKD2.

**Figure 9.**
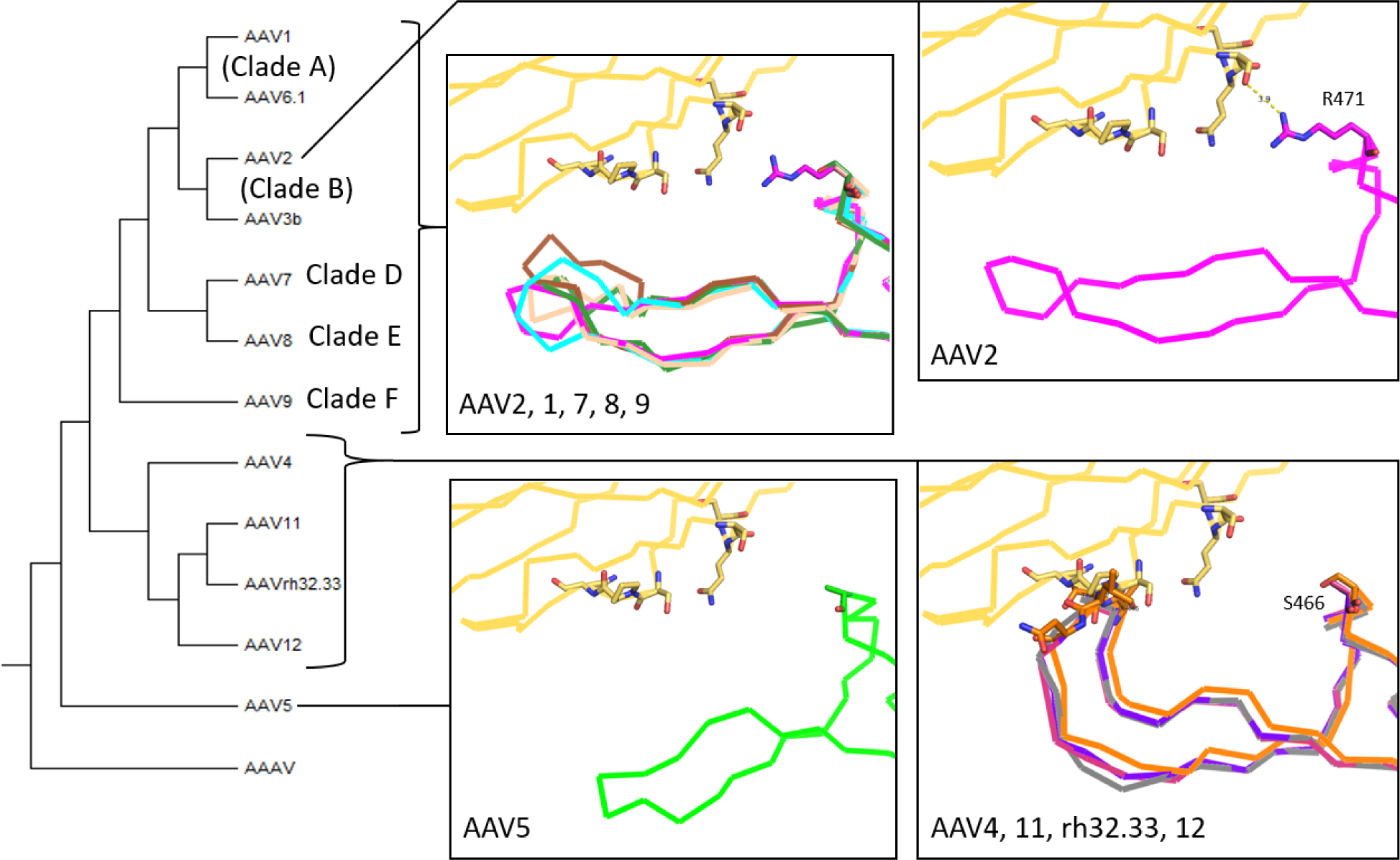
Potential interactions of variable region 4 (VR-IV) for AAV structures after alignment to the AAV2-PKD15 receptor complex (PDBid: 6nz0). Structures and serotype color coding are as described in Figure 4, and AAVR is shown with yellow carbons. A strong steric hindrance was observed between PKD2 (S413, P414 and Q415) and VR-IV for all of the AAV4-clade members (specifically for AAV4, T446 and T447). The phylogenetic tree is based on VP1 sequences.

#### 3.4.2 Comparison of AAV4 with the PKD1-binding site as seen in the AAV5 complex

By contrast to clade B and most other serotypes, the interactions of AAV5 with AAVR are mediated exclusively through PKD1 [27,42]. Even though PKD1 plays an accessory role in the transduction of many serotypes [27], it has been observed, to date, only in the structures of AAV5 complexes [30,31] to which we turn in examining why the AAV4 structure might be incompatible with PKD1-binding. For AAV5, PKD1-interacting residues come from VR-IV, -VII and VR-IX (Table 2). The structures of AAV4 and related clade members were overlaid on the 2.5 Å structure of the AAV5-AAVR complex to rationalize why PKD1-binding is not possible for the AAV4 clade and why avidity is sharply reduced in other non-AAV5 clades.

**Table 2.**
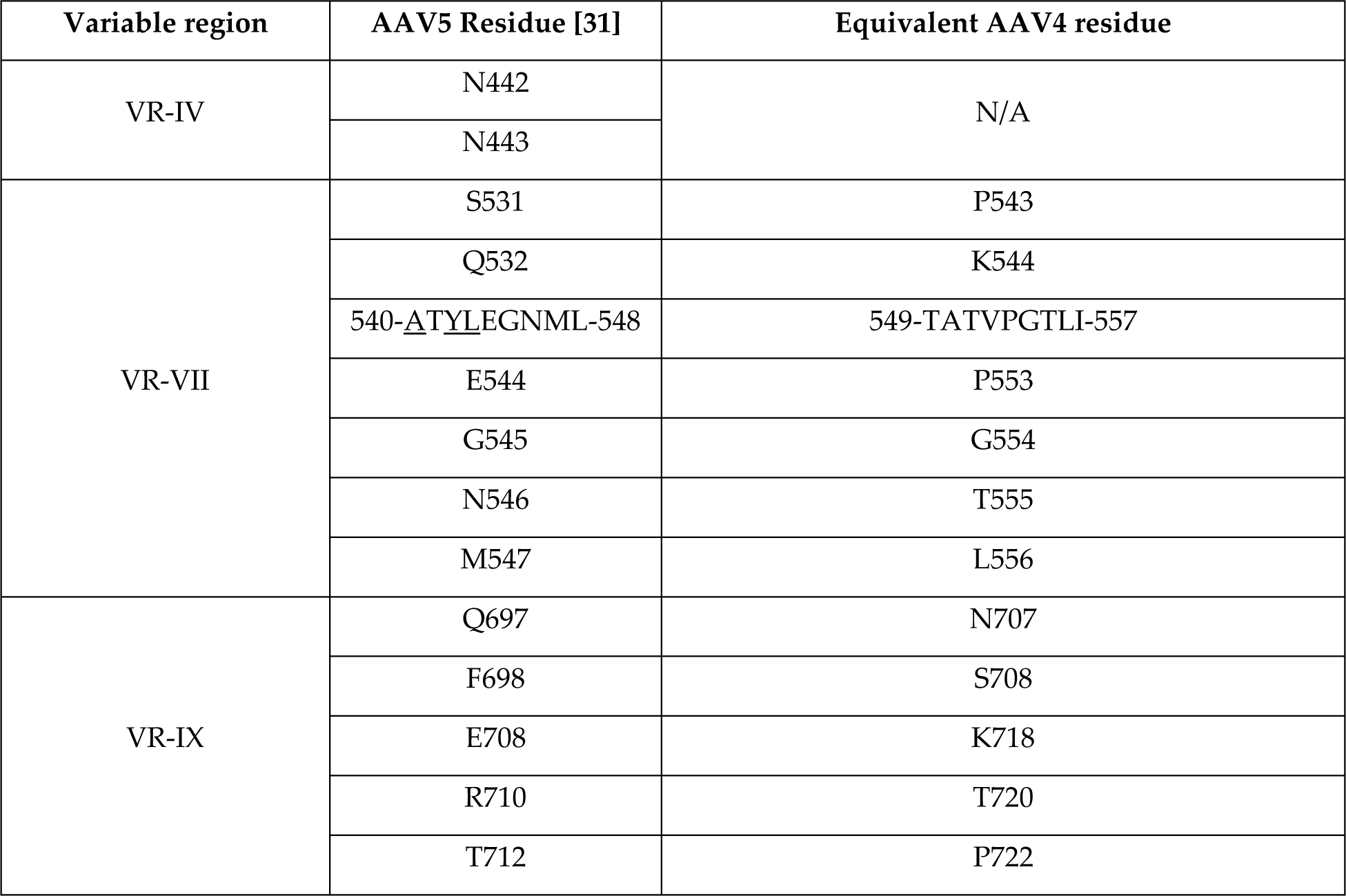
List of contact residues of AAV5 interacting with AAVR [31] and the corresponding residues of AAV4.

| Variable region | AAV5 Residue [31] | Equivalent AAV4 residue |
| --- | --- | --- |
| VR-IV | N442 | N/A |
|  | N443 |  |
| VR-VII | S531 | P543 |
|  | Q532 | K544 |
|  | 540-<u>A</u>T<u>Y</u>L<u>E</u>G<u>N</u>M<u>L</u>-548 | 549-TATVP<u>G</u>T<u>L</u>I-557 |
|  | E544 | P553 |
|  | G545 | G554 |
|  | N546 | T555 |
|  | M547 | L556 |
| VR-IX | Q697 | N707 |
|  | F698 | S708 |
|  | E708 | K718 |
|  | R710 | T720 |
|  | T712 | P722 |

The first region of interest is variable region 4 (VR-IV), with two AAV5 residues contacting PKD1 (Figure 10). VR-IV differs in length between the major branches of the AAV family. The PKD1-binding AAV5 has the shortest loop. AAV4-like viruses have the longest loop, but the additional residues extend the loop away from AAVR-K399. The AAV4-like loop passes K399 with small polar residues – there is no unfavorable steric or electrostatic interaction. By contrast, when aligned to the AAV5 complex, an arginine in AAV2 (R459) or a lysine in AAV9 (K462) is predicted to be in close proximity with AAVR-K399, potentially leading to unfavorable steric and/or electrostatic interactions.

**Figure 10.**
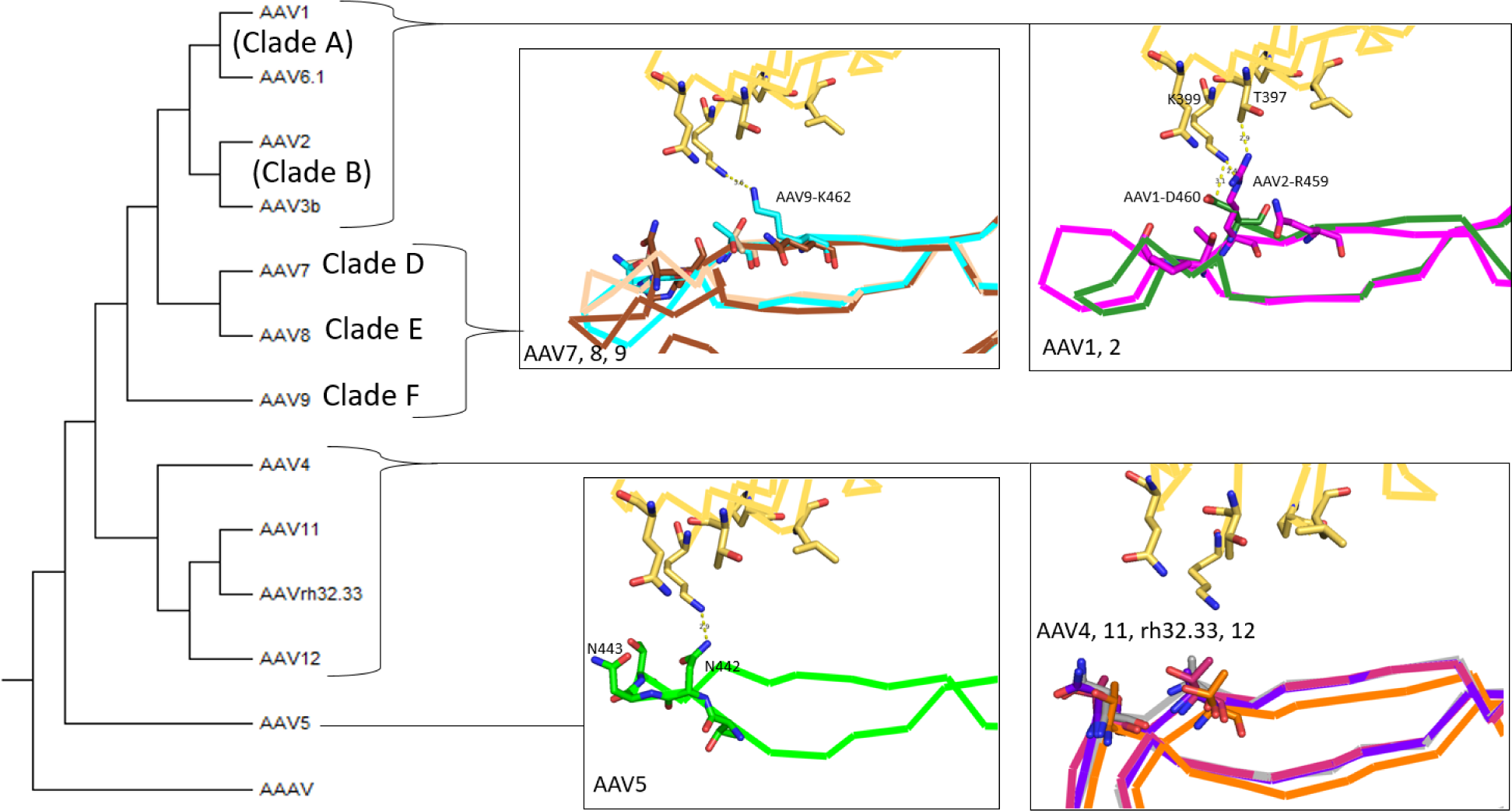
Potential interactions of variable region 4 (VR-IV) for AAV structures after alignment to the AAV5-PKD12 receptor complex (PDBid: 7kpn Structures and serotype color coding are as described in Figure 4, and AAVR is shown with yellow carbons. A steric hindrance, in addition to an electrostatic repulsion, is likely between AAVR-K399 (yellow) and AAV2-R459 (magenta) or AAV9-K462 (cyan). The phylogenetic tree is based on VP1 sequences.

Consider now VR-VII. The AAV5 loop is 3 residues longer than most serotypes and forms a hairpin structure with ample separation from PKD1 (Figure 11). In AAV5, this region does not change significantly whether bound or unbound to AAVR [31]. VR-VII is folded somewhat differently in the AAV4 clade and there would be extensive clashes with a bound PKD1. The most severe would be between AAVR-Y355 and AAV4-K544. For other AAV4-clade members (AAV11, 12 and rh32.33), there are extensive clashes as well. Other clades (e.g., AAV1, 2, 7, 8, 9) superimposed on the AAV5 complex, are predicted to have multiple clashes with AAVR G375-L376 and I349-H351. It appears that all serotypes, except those similar to AAV5, have VR-VII loops that would be completely incompatible with PKD1-binding unless there is an induced conformational change.

**Figure 11.**
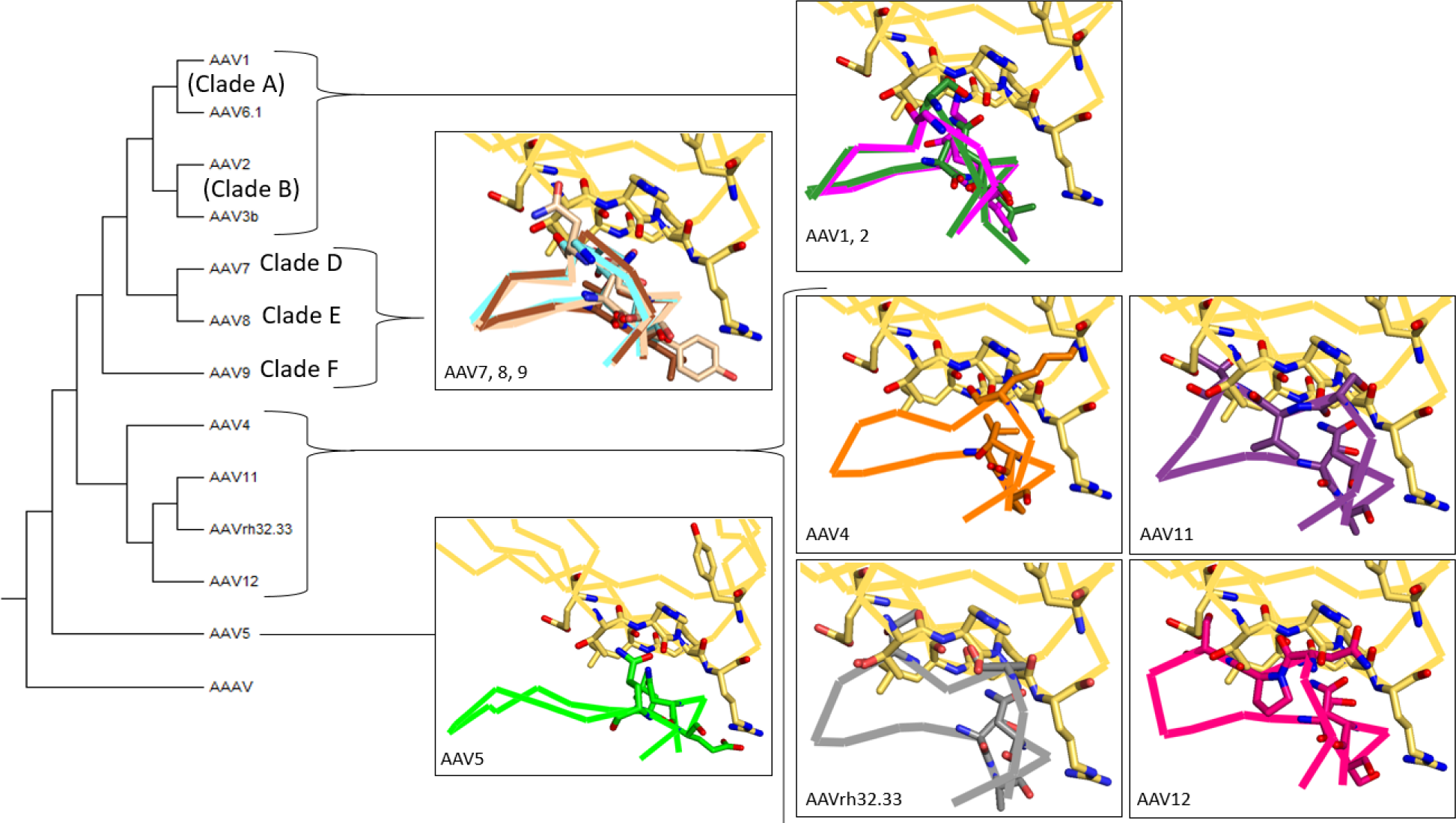
Potential interactions of variable region 7 (VR-VII) for AAV structures after alignment to the AAV5-PKD12 receptor complex (PDBid: 7kpn). Structures and serotype color coding are as described in Figure 4, and AAVR is shown with yellow carbons. A steric clash is observed between AAV4-K544 and AAVR-Y355 and multiple conflicts for the other AAV4-clade members. For serotypes other than AAV5, this region contains multiple contacts that would be incompatible to forming a complex with PKD1. The phylogenetic tree is based on VP1 sequences.

Finally, consider VR-IX (*Figure 12*). The structure of the AAV5 complex shows a favorable electrostatic salt bridge at 2.9 Å between AAVR-H351 and AAV5-E708, which is not conserved. By contrast, the AAV4-clade members have positively charged residues at this location (AAV4-K718; AAV12-H728) that would not interact favorably with AAVR-H351. No other clashes were seen with the other AAV serotypes analyzed.

**Figure 12.**
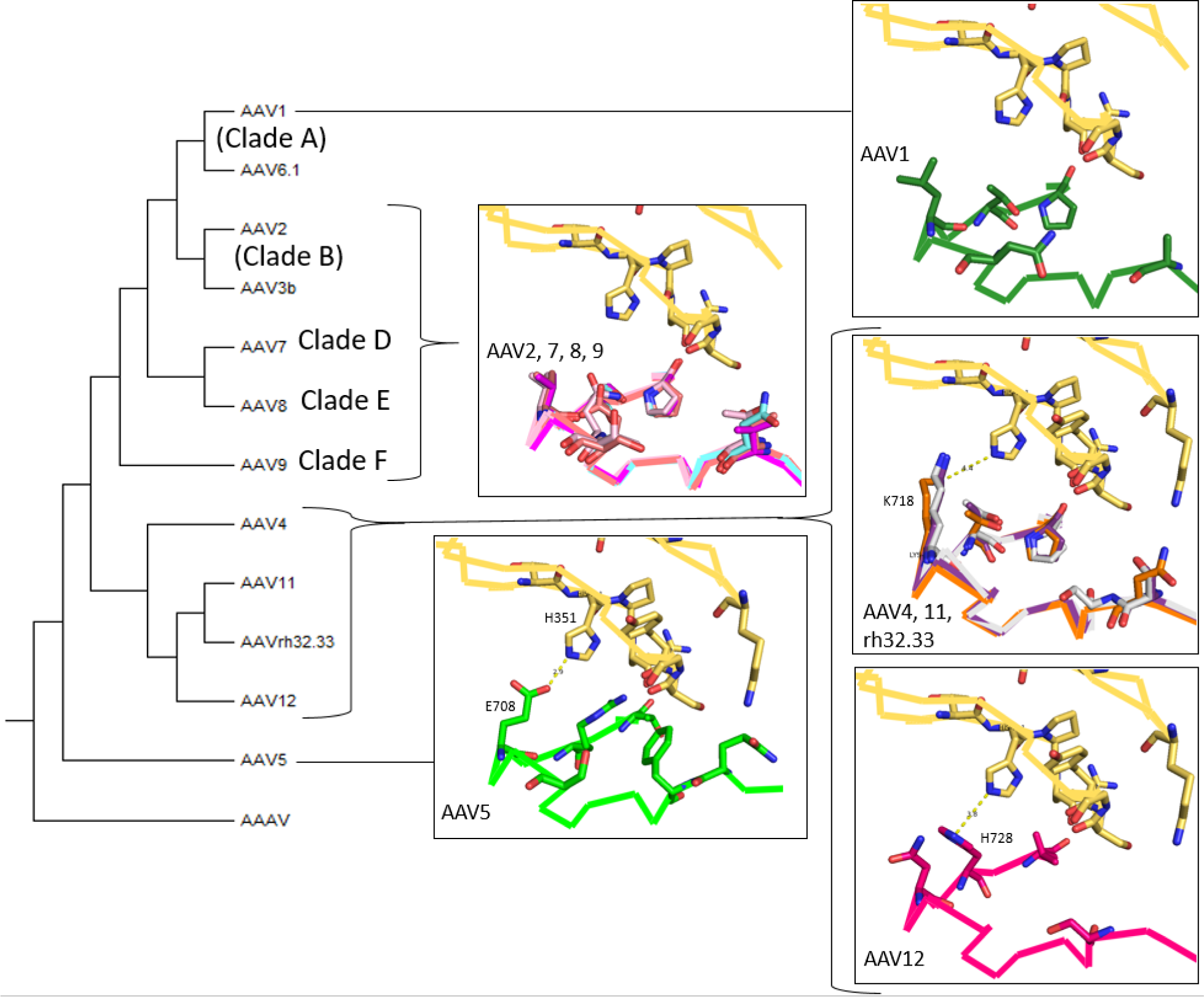
Potential interactions of variable region 9 (VR-IX) for AAV structures after alignment to the AAV5-PKD12 receptor complex (PDBid: 7kpn). Structures and serotype color coding are as described in Figure 4, and AAVR is shown with yellow carbons. No steric hindrance was observed for this region between PKD1 and these serotypes. However, there would likely be unfavorable electrostatics for the AAV4-clade (between AAVR-H351 and AAV4-K718 or AAV12-H728) replacing the salt bridge in AAV5 (E708). The phylogenetic tree is based on VP1 sequences.

## 4. Conclusions

In this article, we report a high-resolution structure of AAV4 at 2.21 Å. We then compared it with AAV structures known to interact with AAVR to suggest reasons why AAV4 might not be able to form a complex with the receptor, which is necessary for all other AAV serotypes to infect cells. We specifically considered the PKD2 and PKD1 portions of AAVR that are known to interact with AAV2 and AAV5, respectively.

Multiple differences were identified between AAV4 and AAV2 which may explain why binding might not be possible between AAV4 and PKD2 (see Figure 13 for a summary). Many of the residues identified as contact residues in AAV2 are different in the AAV4 clade. Regions where differences might be of highest impact include VR-I, -III, -IV, and -V, which result in multiple steric clashes for a potential interaction with PKD2. AAV5, like AAV4, is PKD2-incompatible. AAV5 VR-I and -III would be in conflict with AAV2-like binding of PKD2, but AAV5 is more similar to PKD2-binding serotypes in other loops of AAV2’s binding site.

**Figure 13.**
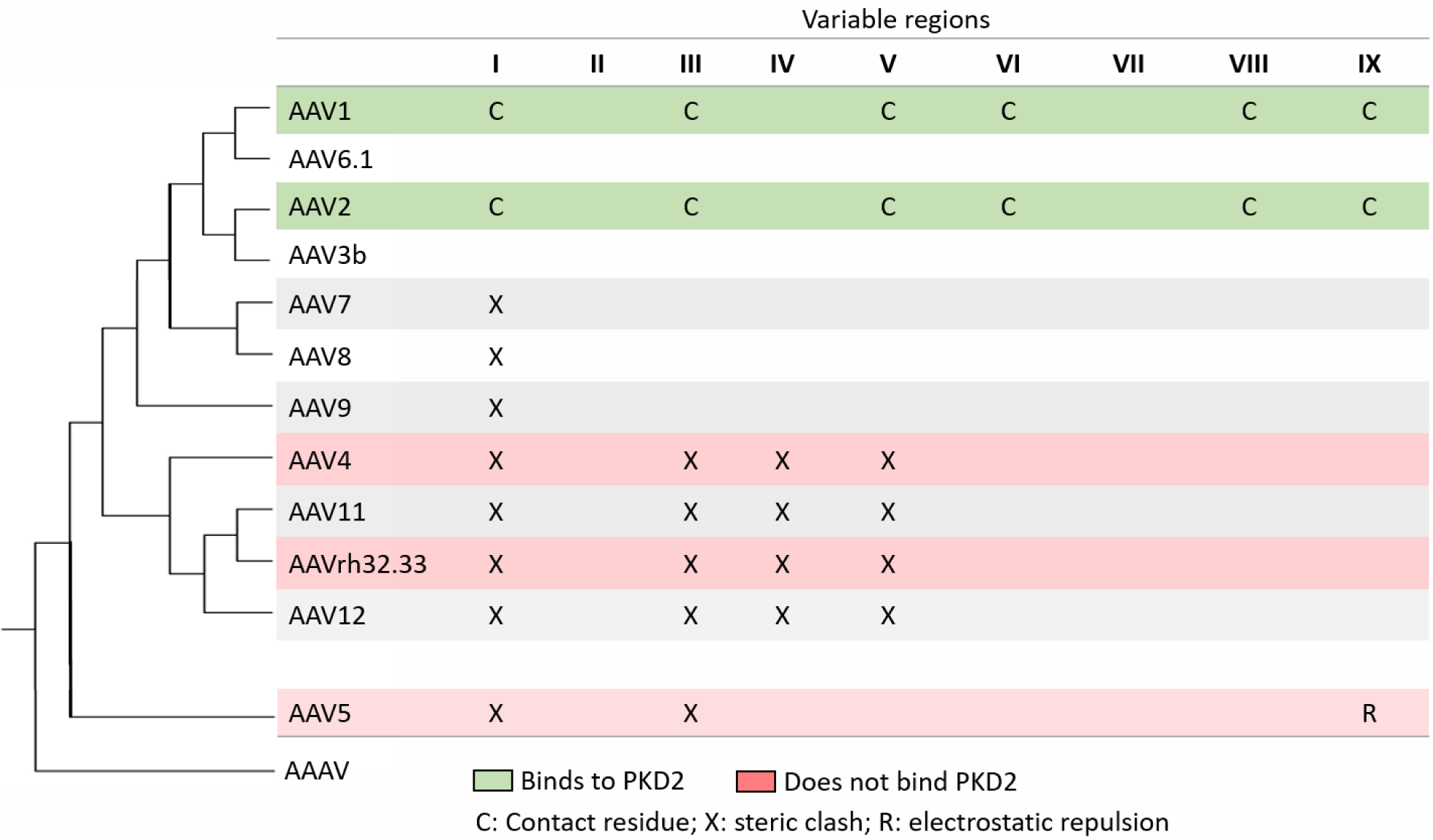
Summary of interactions between PKD2 and various serotypes.

In terms of understanding why AAV4 and other clades does not bind PKD1 like AAV5, the answer appears simpler (Figure 14). It is AAV5 alone that has a VR-VII conformation that is compatible with binding. It is possible that differences in VR-IX could have a secondary role. We see a favorable salt bridge in the AAV5 complex lost in other clades, and with a potentially unfavorable electrostatic environment for H351 in the AAV4 clade.

**Figure 14.**
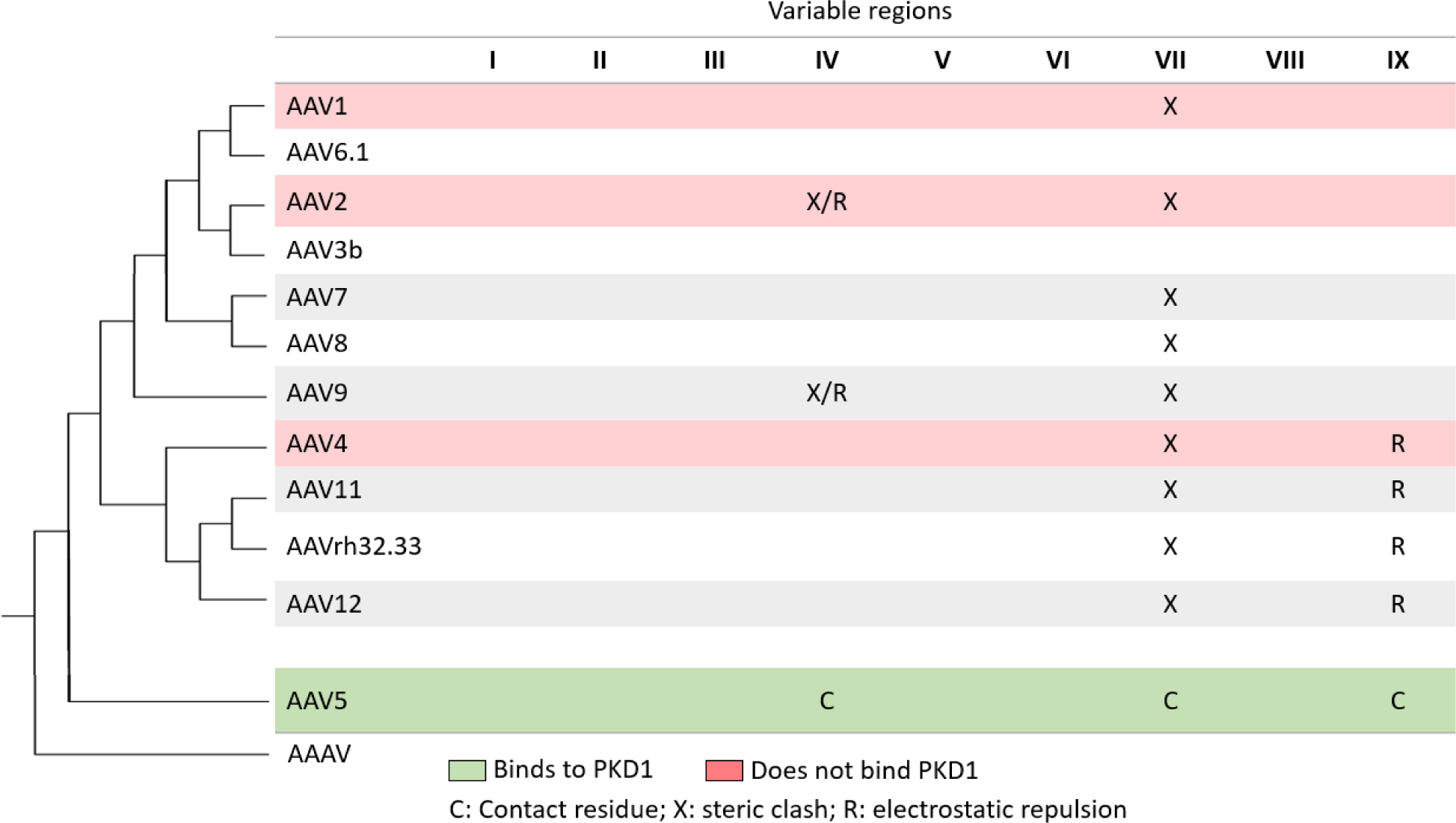
Summary of predicted interactions between PKD1 and various serotypes.

With on-going interest in AAV as the delivery vector of gene therapy treatments, research into understanding the mechanisms of virus-receptor interactions is an essential foundation for improving the efficiency and specificity of gene delivery. The structure of AAV4 presented here, and the subsequent comparative analysis using receptor complexes of homologous strains, has allowed a narrowing down of the viral determinants of the modes of binding to the predominant cellular receptor. There appears to be only one conformation of surface loop VR-VII that is compatible with AAV binding to the PKD1 domain of AAVR, like AAV5. For AAV2-like binding to the PKD2 domain of AAVR, several loops need to be configured appropriately: VR-III, VR-IV, VR-V and VR-I. These are located in two regions of the viral surface: VR-III and VR-I are on the shoulder extending down from each spike towards the 2-fold axis; and VR-IV and VR-V are further towards the top of the spikes (Figure 2). These two regions interact respectively with the N-terminal and C-terminal ends of the PKD2 domain, as tightly bound in the AAV2 complex (Figure 3).

## Author Contributions

Conceptualization, M.S.C.; validation, M.A.S. and G.M.Z.; formal analysis, G.M.Z., M.A.S. and M.S.C.; investigation, G.M.Z. and M.A.S.; data collection, N.L.M.; data curation, G.M.Z. and M.A.S.; writing—original draft preparation, G.M.Z. and M.S.C.; writing—review and editing, G.M.Z., M.A.S. and M.S.C.; visualization, G.M.Z.; supervision, M.S.C. and T.W.; project administration, M.S.C.; funding acquisition, M.S.C. All authors have read and agreed to the published version of the manuscript.

## Funding

This research was funded by a grant from The National Institute of Health (NIH), R35 GM122564. High resolution EM data were collected performed at the Pacific Northwest Center for CryoEM at Oregon Health Sciences University, supported by NIH grant, U24GM129547, and applied for through EMSL (grid.436923.9), a DOE Office of Science User Facility sponsored by the Office of Biological and Environmental Research.

## Acknowledgments

Special thanks to Mizzou Microscopy core and the Pacific Northwest Center for Cryo-EM for their role in data collection. Thanks to the USDA, ARS, BCIRL, lab in Columbia, MO for supplying a starter culture of Sf9 cells used to produce protein. Much gratitude goes to Dr. Kyle Stiers and the Beamer Lab for assistance with the Thermal Shift Assays and access to the custom software for analysis. The AAV4 cryo-EM maps will be made available from the EMDB with accession number 25903. The AAV4 atomic coordinates will be available from the PDB data base as 7thr.

## Conflicts of Interest

The authors declare no conflict of interest.

## Notes

### Competing Interest Statement

The authors have declared no competing interest.

## References

1. Naso, M.F. et al. (2017) Adeno-Associated Virus (AAV) as a Vector for Gene Therapy. BioDrugs 31, 317–334. 10.1007/s40259-017-0234-5

2. Mendell, J.R. et al. (2021) Current Clinical Applications of In Vivo Gene Therapy with AAVs. Mol Ther 29, 464–488. 10.1016/j.ymthe.2020.12.007

3. Hoy, S.M. (2019) Onasemnogene Abeparvovec: First Global Approval. Drugs 79, 1255–1262. 10.1007/s40265-019-01162-5

4. de Jong, Y.P. and Herzog, R.W. (2021) Liver gene therapy and hepatocellular carcinoma: A complex web. Mol Ther 29, 1353–1354. 10.1016/j.ymthe.2021.03.009

5. Dalwadi, D.A. et al. (2021) Liver Injury Increases the Incidence of HCC following AAV Gene Therapy in Mice. Mol Ther 29, 680–690. 10.1016/j.ymthe.2020.10.018

6. Morales, L. et al. (2020) Broader Implications of Progressive Liver Dysfunction and Lethal Sepsis in Two Boys following Systemic High-Dose AAV. Mol Ther 28, 1753–1755. 10.1016/j.ymthe.2020.07.009

7. Philippidis, A. (2020) After Third Death, Audentes’ AT132 Remains on Clinical Hold. Hum Gene Ther 31, 908–910. 10.1089/hum.2020.29133.bfs

8. Muzyczka, N. and Berns, K.I. (2001) Parvoviridae: The Viruses and Their Replication. In Virology (4 edn) (Fields, B.N.et al., eds), pp. 2327–2360, Lippincott Williams & Wilkins

9. Bleker, S. et al. (2005) Mutational analysis of narrow pores at the fivefold symmetry axes of adeno-associated virus type 2 capsids reveals a dual role in genome packaging and activation of phospholipase A2 activity. J Virol 79, 2528–2540. 10.1128/JVI.79.4.2528-2540.2005

10. Girod, A. et al. (2002) The VP1 capsid protein of adeno-associated virus type 2 is carrying a phospholipase A2 domain required for virus infectivity. J Gen Virol 83, 973–978

11. Popa-Wagner, R. et al. (2012) Impact of VP1-specific protein sequence motifs on adeno-associated virus type 2 intracellular trafficking and nuclear entry. J Virol 86, 9163–9174. 10.1128/JVI.00282-12

12. Kurian, J.J. et al. (2019) Adeno-Associated Virus VP1u Exhibits Protease Activity. Viruses 11. 10.3390/v11050399

13. Johnson, J.S. et al. (2010) Mutagenesis of adeno-associated virus type 2 capsid protein VP1 uncovers new roles for basic amino acids in trafficking and cell-specific transduction. J Virol 84, 8888–8902. 10.1128/JVI.00687-10

14. Kronenberg, S. et al. (2001) Electron cryo-microscopy and image reconstruction of adeno-associated virus type 2 empty capsids. EMBO Rep 2, 997–1002

15. Kronenberg, S. et al. (2005) A conformational change in the adeno-associated virus type 2 capsid leads to the exposure of hidden VP1 N termini. J Virol 79, 5296–5303

16. Gerlach, B. et al. (2011) Conformational changes in adeno-associated virus type 1 induced by genome packaging. J Mol Biol 409, 427–438. 10.1016/j.jmb.2011.03.062

17. Hu, G. et al. (2022) Adeno-associated Virus Receptor-binding: Flexible Domains and Alternative Conformations through cryo-Electron Tomography of AAV2 and AAV5 complexes. bioRxiv, 2022.2001.2010.475736. 10.1101/2022.01.10.475736

18. Sonntag, F. et al. (2006) Adeno-associated virus type 2 capsids with externalized VP1/VP2 trafficking domains are generated prior to passage through the cytoplasm and are maintained until uncoating occurs in the nucleus. J Virol 80, 11040–11054

19. Lins-Austin, B. et al. (2020) Adeno-Associated Virus (AAV) Capsid Stability and Liposome Remodeling During Endo/Lysosomal pH Trafficking. Viruses 12. 10.3390/v12060668

20. Venkatakrishnan, B. et al. (2013) Structure and Dynamics of Adeno-Associated Virus Serotype 1 VP1-Unique N-Terminal Domain and Its Role in Capsid Trafficking. J Virol 87, 4974–4984. 10.1128/JVI.02524-12

21. Madigan, V.J. et al. (2020) The Golgi Calcium ATPase Pump Plays an Essential Role in Adeno-associated Virus Trafficking and Transduction. J Virol 94. 10.1128/JVI.01604-20

22. Summerford, C. and Samulski, R.J. (1998) Membrane-associated heparan sulfate proteoglycan is a receptor for adeno-associated virus type 2 virions. J Virol 72, 1438–1445

23. Walters, R.W. et al. (2001) Binding of adeno-associated virus type 5 to 2,3-linked sialic acid is required for gene transfer. J Biol Chem 276, 20610–20616. 10.1074/jbc.M101559200

24. Pillay, S. et al. (2016) An essential receptor for adeno-associated virus infection. Nature 530, 108–112. 10.1038/nature16465

25. Dudek, A.M. et al. (2020) GPR108 Is a Highly Conserved AAV Entry Factor. Mol Ther 28, 367–381. 10.1016/j.ymthe.2019.11.005

26. Poon, M.W. et al. (2011) Dyslexia-associated kiaa0319-like protein interacts with axon guidance receptor nogo receptor 1. Cell Mol Neurobiol 31, 27–35. 10.1007/s10571-010-9549-1

27. Pillay, S. et al. (2017) Adeno-associated Virus (AAV) Serotypes Have Distinctive Interactions with Domains of the Cellular AAV Receptor. J Virol 91, e00391–00317. 10.1128/JVI.00391-17

28. Meyer, N.L. et al. (2019) Structure of the gene therapy vector, adeno-associated virus with its cell receptor, AAVR. Elife 8, e44707. 10.7554/eLife.44707

29. Zhang, R. et al. (2019) Adeno-associated virus 2 bound to its cellular receptor AAVR. Nature Microbiology 4, 675–682. 10.1038/s41564-018-0356-7

30. Zhang, R. et al. (2019) Divergent engagements between adeno-associated viruses with their cellular receptor AAVR. Nat Commun 10, 3760. 10.1038/s41467-019-11668-x

31. Silveria, M.A. et al. (2020) The Structure of an AAV5-AAVR Complex at 2.5 A Resolution: Implications for Cellular Entry and Immune Neutralization of AAV Gene Therapy Vectors. Viruses 12, 1326. 10.3390/v12111326

32. Govindasamy, L. et al. (2006) Structurally mapping the diverse phenotype of adeno-associated virus serotype 4. J Virol 80, 11556–11570. 10.1128/JVI.01536-06

33. Xie, Q. et al. (2002) The atomic structure of adeno-associated virus (AAV-2), a vector for human gene therapy. Proc Natl Acad Sci U S A 99, 10405–10410. 10.1073/pnas.162250899

34. Padron, E. et al. (2005) Structure of adeno-associated virus type 4. J Virol 79, 5047–5058

35. Cheng, Y. (2015) Single-Particle Cryo-EM at Crystallographic Resolution. Cell 161, 450–457. 10.1016/j.cell.2015.03.049

36. Scheres, S.H. (2016) Processing of Structurally Heterogeneous Cryo-EM Data in RELION. Methods Enzymol 579, 125–157. 10.1016/bs.mie.2016.04.012

37. Baldwin, P.R. et al. (2018) Big data in cryoEM: automated collection, processing and accessibility of EM data. Curr Opin Microbiol 43, 1–8. 10.1016/j.mib.2017.10.005

38. Tan, Y.Z. et al. (2018) Sub-2 Å Ewald curvature corrected structure of an AAV2 capsid variant. Nature Communications 9, 3628. 10.1038/s41467-018-06076-6

39. Xie, Q. et al. (2020) Adeno-Associated Virus (AAV-DJ)-Cryo-EM Structure at 1.56 A Resolution. Viruses 12, 1194. 10.3390/v12101194

40. Mietzsch, M. et al. (2021) Completion of the AAV Structural Atlas: Serotype Capsid Structures Reveals Clade-Specific Features. Viruses 13. 10.3390/v13010101

41. Mikals, K. et al. (2014) The structure of AAVrh32.33, a novel gene delivery vector. J Struct Biol 186, 308–317. 10.1016/j.jsb.2014.03.020

42. Dudek, A.M. et al. (2018) An Alternate Route for Adeno-associated Virus (AAV) Entry Independent of AAV Receptor. J Virol 92. 10.1128/JVI.02213-17

43. Li, M.Z. and Elledge, S.J. (2007) Harnessing homologous recombination in vitro to generate recombinant DNA via SLIC. Nat Methods 4, 251–256. 10.1038/nmeth1010

44. Urabe, M. et al. (2002) Insect cells as a factory to produce adeno-associated virus type 2 vectors. Hum Gene Ther 13, 1935–1943

45. Meyer, N. et al. (2020) Expression and Purification of Adeno-associated Virus Virus-like Particles in a Baculovirus System and AAVR Ectodomain Constructs in E. coli. Bio-Protocol 10, e3513. 10.21769/BioProtoc.3513

46. Scheres, S.H. (2012) A Bayesian view on cryo-EM structure determination. J Mol Biol 415, 406–418. 10.1016/j.jmb.2011.11.010

47. Rohou, A. and Grigorieff, N. (2015) CTFFIND4: Fast and accurate defocus estimation from electron micrographs. J Struct Biol 192, 216–221. 10.1016/j.jsb.2015.08.008

48. Chapman, M.S. et al. (2013) Atomic modeling of cryo-electron microscopy reconstructions--joint refinement of model and imaging parameters. J Struct Biol 182, 10–21. 10.1016/j.jsb.2013.01.003

49. Emsley, P. et al. (2010) Features and development of Coot. Acta Crystallogr D Biol Crystallogr 66, 486–501. 10.1107/S0907444910007493

50. Brünger, A.T. et al. (1998) Crystallography and NMR system: A new software system for macromolecular structure determination. Acta Crystallographica D54, 905–921

51. Andreotti, G., M. Monticelli and V. Cubellis (2015) Looking for protein stabilizing drugs with thermal shift assay. Drug testing and analysis 7, 4. DOI 10.1002/dta.1798

52. Nam, H.J. et al. (2007) Structure of adeno-associated virus serotype 8, a gene therapy vector. J Virol 81, 12260–12271. 10.1128/JVI.01304-07

53. DiMattia, M.A. et al. (2012) Structural insight into the unique properties of adeno-associated virus serotype 9. J Virol 86, 6947–6958. 10.1128/JVI.07232-11

54. Larkin, M.A., G. Blackshields, N.P. Brown, R. Chenna, P.A. McGettigan, H. McWilliam, F. Valentin, I.M. Wallace, A. Wilm, R. Lopez, J.D. Thompson, T.J. Gibson, D.G. Higgins (2007) Clustal W and Clustal X version 2.0. Bioinformatics 23, 2. https://doi.org/10.1093/bioinformatics/btm404

55. DeLano, W.L. (2002). The PyMOL Molecular Graphics System. DeLano Scientific

56. (2015). The PyMOL Molecular Graphics System. 2.0 ed. Schrödinger, LLC.

57. Rieser, R. et al. (2020) Intrinsic Differential Scanning Fluorimetry for Fast and Easy Identification of Adeno-Associated Virus Serotypes. J Pharm Sci 109, 854–862. 10.1016/j.xphs.2019.10.031

58. Pacouret, S. et al. (2017) AAV-ID: A Rapid and Robust Assay for Batch-to-Batch Consistency Evaluation of AAV Preparations. Mol Ther 25, 1375–1386. 10.1016/j.ymthe.2017.04.001

59. Bennett, A. et al. (2017) Thermal Stability as a Determinant of AAV Serotype Identity. Mol Ther Methods Clin Dev 6, 171–182. 10.1016/j.omtm.2017.07.003

