## Supplementary materials for "Cryo-EM Structure of Adeno-associated virus-4 at 2.2 Å resolution"

*Table S 1: Primers used for cloning of AAV4 into pFastBac-LIC(4A).*

| Name | Sequence[a,b] | Purpose |
|---|---|---|
| GMZ-1b | GACTAGTGAGCTCGTCGACGTAGG | Amplification of pFastBac-LIC(4A); reverse primer |
| GMZ-2 | CACTGATTAAGCATTGGTAACTGTCAGACC | Amplification of pFastBac-LIC(4A); forward primer |
| GMZ-3 | GGTCTGACAGTTACCAATGCTTAATCAGTG | Amplification of pFastBac-LIC(4A); reverse primer |
| GMZ-4 | GTACCAAGCTTGTCGAGAAGTACTAGAGGATC | Amplification of pFastBac-LIC(4A); forward primer |
| GMZ-7b | CCTACGTCGACGAGCTCACTAGTC<br>AcGACTGACGGTTACCTTCCAGATTG | Amplification of the AAV4-VP1 sequence; forward primer |
| GMZ-8 | GATCCTCTAGTACTTCTCGACAAGCTTGGTAC<br>TTACAGGTGGTGGGTGAGGTAGC | Amplification of the AAV4-VP1 sequence; reverse primer |
| GMZ-9 | CGTATACTCCGGACTATTAATAGATCATGGAG | Sequencing primer |
| GMZ-10 | AACCTCTACAAATGTGGTATGGCTG | Sequencing primer |
| EEL23 | CAGTGAGATGCGTGCAGCAG | Sequencing primer |
| EEL24 | ATGCCTTCTACTGCCTGGAG | Sequencing primer |
| EEL25 | TGACCAGAGCAACAGCAACC | Sequencing primer |
| EEL27 | GAAGCCCTGCTGCTTGATTG | Sequencing primer |
| EEL28 | GCCCAAACCCACCAATCAGC | Sequencing primer |

a – underlined sequence reflects the overhang sequence shared with vector sequence.

b – double underlined base shows location of mutation within VP1 start codon.

*Table S 2. Melting temperatures of isolated VLPs for various serotypes and the values published by others to show range of reported values.*

| Serotype | Average Tm ± standard deviation | Riser et al. [57] | Pacouret et al. [58] | Bennett et al. [59] |
|---|---|---|---|---|
| AAV2 | 73.4°C ± 0.13 | 68°C | 69.6°C | 77.5°C |
| AAV5 | 89.9°C ± 0.06 | 90°C | 88.7°C | 89.7°C |
| AAV4 | 79.9°C ± 0.03 | - | - | 76.0°C |

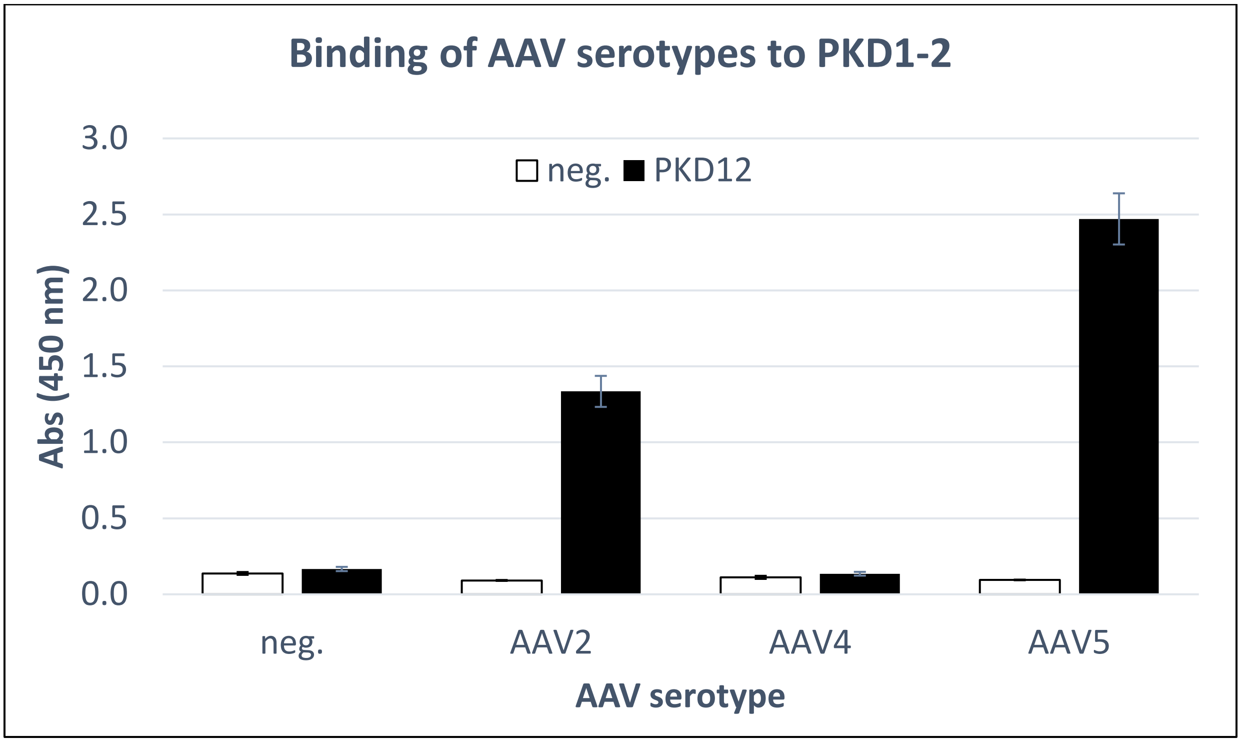


*Figure S 1. ELISA results of PKD1-2 binding to AAV2, AAV4 and AAV5. No added ligand (clear) compared with PKD1-2 bound samples (filled).*

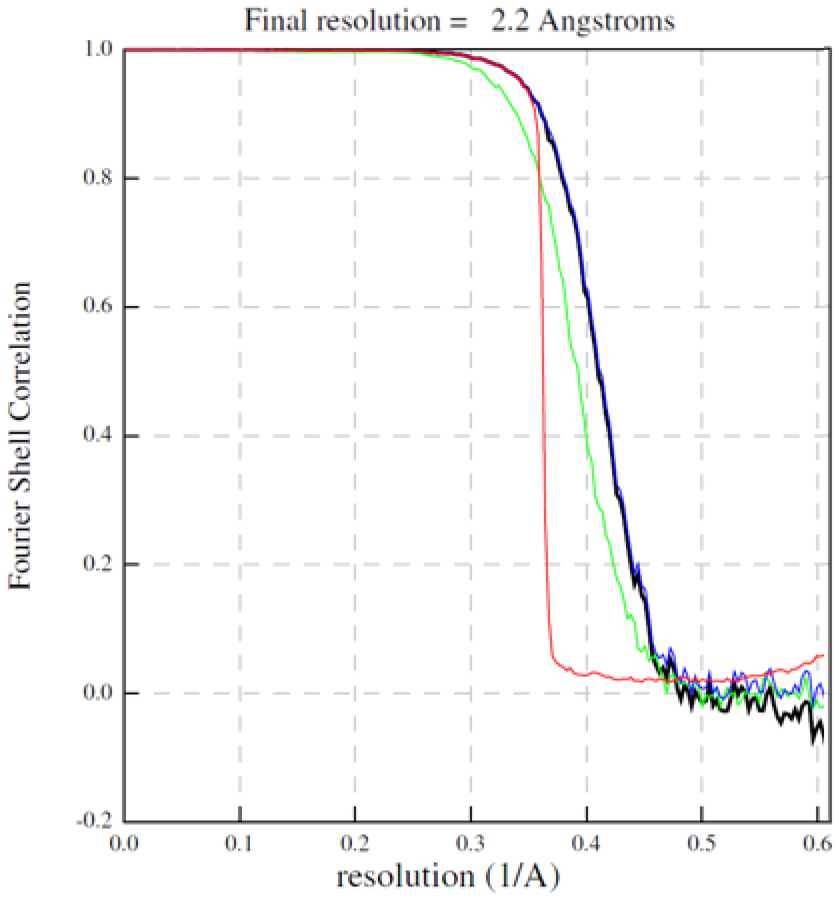


*Figure S 2. FSC curve for AAV4.*

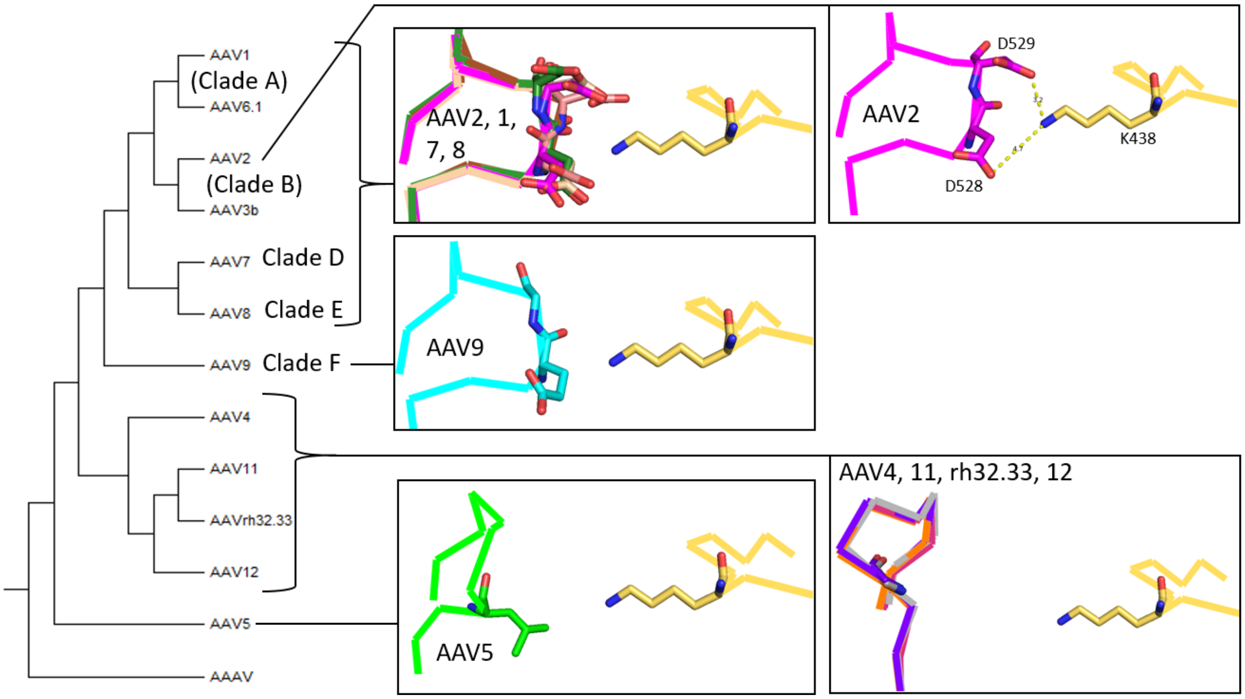


*Figure S 3. Potential interactions of variable region 6 (VR-VI) for AAV structures after alignment to the AAV2-PKD15 receptor complex (PDBid: 6nz0). Structures and serotype color coding are as described in Figure 4, and AAVR is shown with yellow carbons. No steric hindrance was observed for this region between PKD2 and these serotypes. Residues in AAV5 and the AAV4-clade that correspond to contact residues in AAV2 are not as close to AAVR. Additionally, an aspartic acid (AAV2-D529 which is in contact with AAVR-K438) is conserved in clades A-E but is a glycine (G530) in AAV9. The phylogenetic tree is based on VP1 sequences.*

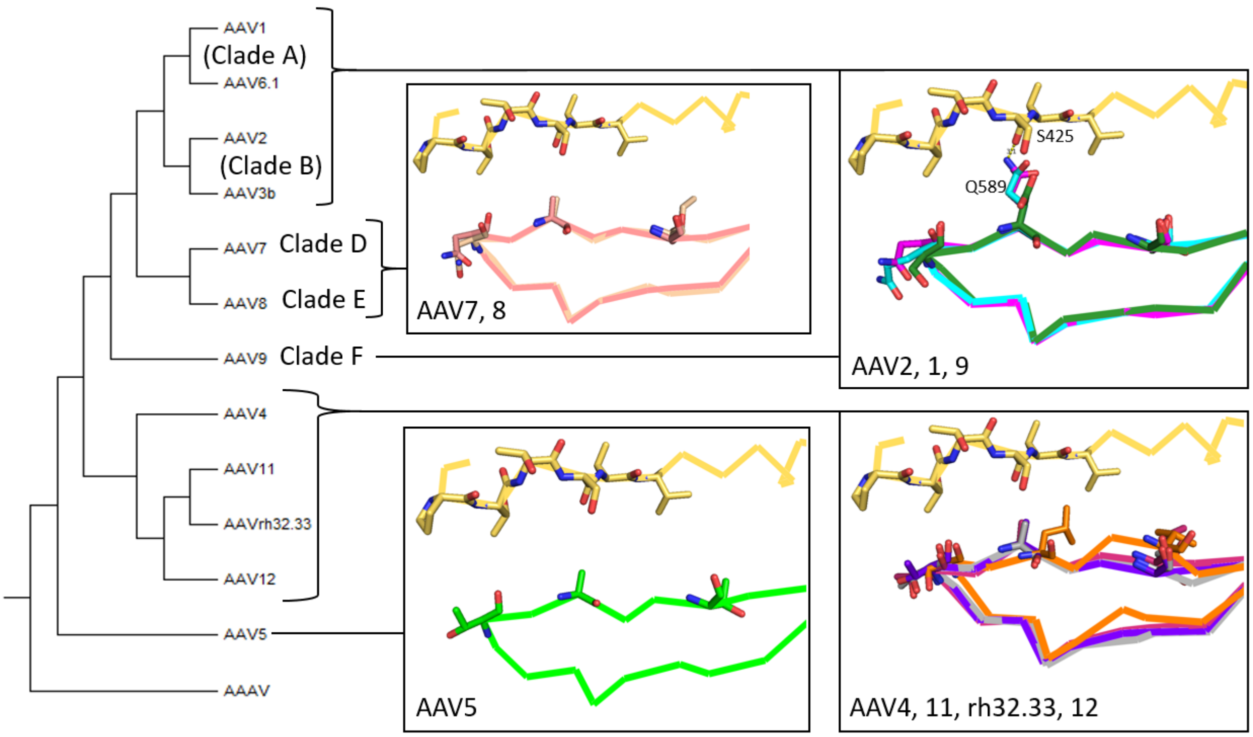


*Figure S 4. Potential interactions of variable region 8 (VR-VIII) for AAV structures after alignment to the AAV2-PKD15 receptor complex (PDBid: 6nz0). Structures and serotype color coding are as described in Figure 4, and AAVR is shown with yellow carbons. No steric hindrance was observed for any of the serotypes. The contact residue AAV2-Q589 (in contact with AAVR-S425) is conserved in serotype 9, an aspartate in AAV1 but an alanine in most other strains. The phylogenetic tree is based on VP1 sequences.*
